# Competitor-induced plasticity modifies the interactions and predicted competitive outcomes between annual plants

**DOI:** 10.1101/2024.06.06.597843

**Authors:** Theo L. Gibbs, Jonathan M. Levine, Martin M. Turcotte

## Abstract

Species can modify their traits in response to changes in the environment – a process known as phenotypic plasticity. Because species traits can plastically respond to competition, the competitive effect of one individual on another involves not only reductions in performance, but also changes in morphology, behavior, phenology or physiology that affect interactions with other individuals. In this context, plasticity is often argued to favor species coexistence by increasing the niche differentiation between species, though experimental support that explicitly considers competitive outcomes is largely lacking. Here, we transiently subjected four annual plant species to early-season intraspecific or interspecific competition to explicitly induce plastic responses, and then examined the response of these individuals to other competitors. By measuring the interactions between the species with and without early-season competitors, we isolated the impact of plasticity on species interactions and coexistence. Growing with nearby competitors early in life impacted plant traits including height and morphology. These plastic responses tended to amplify the sensitivity of individuals to competition, and particularly so for interspecific competition. This increase in inter-relative to intraspecific competition caused plasticity to decrease the predicted likelihood of pairwise coexistence. By combining recent theory with a new experimental approach, we provide a pathway towards integrating phenotypic plasticity into our quantitative understanding of coexistence.

## Introduction

Species exhibit remarkable intraspecific variation in traits [1, 2, 3, 4, 5]. One important cause of this variation is phenotypic plasticity, in which the traits of an organism change in response to its environment [6]. Phenotypic plasticity has been observed across taxa and scales of biological organization – plant species adjust their rooting architecture to preferentially grow towards resources [7, 8, 9, 10, 11, 12, 13, 14], many bacterial strains regulate the uptake rates of nutrients depending on which other resources are available [15, 16, 17, 18, 19, 20, 21] and aquatic predators adjust their diets in response to competition [22, 23, 24]. In all three of these examples, plasticity does not just modify the traits of a given species, but also affects the interactions between species in a community.

These examples illuminate a potential feedback, mediated via plastcitiy, between species interactions and traits [25, 26]. When species compete, they not only affect the performance of their competitors by drawing down shared limiting resources, but can also induce plastic changes that make their competitors more (or less) susceptible to other competitors. These plastic changes can then feed back to shape the trajectory of resource competition. Nonetheless, empirical evidence for how plastic shifts in response to competitors shape the competitive outcomes between species is poorly resolved, because phenotypic plasticity and coexistence are usually studied separately [25, 27, 28, 29, 30, 31] (but see [32]).

Phenotypic plasticity research has demonstrated that competitors can induce changes in the traits of a focal individual [22, 24, 33, 34, 35, 36], but how these changes affect competitive outcomes is rarely shown. Instead, these trait changes are often *a priori* interpreted as evidence for increased niche differentiation [31]. However, inferences about coexistence outcomes require measurements of species interactions, rather than just trait measurements, because only these interactions directly predict species’ eventual abundances [37]. Moreover, the relationship between traits and interactions is hardly straightforward [38, 39], so it is not possible to predict interactions from traits alone, except in simple cases [40]. Even when previous work has measured how plasticity shifts interaction strengths, it has focused on changes in interspecific interaction strengths [31]. But it is the relationship between interspecific and intraspecific interactions that dictates whether coexistence is possible, so if plasticity also modifies intraspecific interactions significantly, prior conclusions about how plasticity changes coexistence may be altered.

Coexistence research, by contrast, has successfully quantified the interactions between species, and then used these interactions to predict coexistence outcomes [37, 41]. But standard approaches to understanding species coexistence often assume that species’ traits, and hence their per capita competitive effects, are fixed regardless of their neighbors’ identity or abundance. When phenotypic plasticity modifies the interactions between competing species, the strength of the interactions themselves will depend on the competitive environment, causing the standard tools of coexistence theory to not readily apply. As a result, we lack clear theoretical expectations for how the coexistence between competitors changes when their per capita interactions vary with species density.

Fortunately, there has been recent progress on a very similar theoretical problem where trait (and hence interaction) changes come about from rapid evolution [42]. The key insight from Yamamichi et al. [42] is that coexistence theory metrics remain valid even when interactions are varying as long as they are measured under the appropriate competitive contexts. For example, when determining whether or not one species can invade another, the competitive effect of the resident on the invader, as well as the effect of the resident on itself, must be evaluated assuming that both species have evolved to the common resident. From these invasion growth rates, it is possible to compute an eco-evolutionary niche difference, which can in turn be decomposed into contributions reminiscent of character displacement [43, 44, 45] and the evolution of competitive ability [46, 47, 48, 49, 50, 51]. Interestingly, this recent theory applies to any density-mediated changes in interaction strengths, not just those caused by rapid evolution, and so can readily be adapted to understand how phenotypic plasticity impacts species coexistence.

Here, we leverage this theoretical advance to predict competitive outcomes using experiments that measure the effect of phenotypic plasticity on the interactions between four species of annual plant. In this work, we use an operational definition of phenotypic plasticity that includes any phenotypic change that results from induction by competitors. This definition is broad enough to encompass passive and active plasticity [52, 53, 54, 55, 29], and chosen to include all forms that affect interactions. Inspired by the foundational approach of Relyea (2002) [56], we expose species to either intraspecific or interspecific competitors early in the growing season to experimentally induce plastic responses, and then remove these competitors to minimize their direct competitive impact (Fig. 1A). We then measure the interactions between species both with and without early-season competitors to quantify how plastic changes impact interaction strengths (Fig. 1B). Importantly, measuring the interactions between species when they have and have not plastically responded to early-season competitors allows us to disentangle the direct competitive effect of inducers from their modification of subsequent interaction strengths [31]. Moreover, our experimental treatments allow us to parameterize the theoretical framework developed by Yamamichi et al. [42], and hence determine how plasticity impacts coexistence outcomes. Specifically, we address three main questions: 1) How does phenotypic plasticity affect the strength of intraspecific and interspecific competitive interactions? 2) What are the consequences of plasticity for species coexistence and niche differentiation? 3) Do competitors induce changes in the traits of focal species?

**Figure 1:**
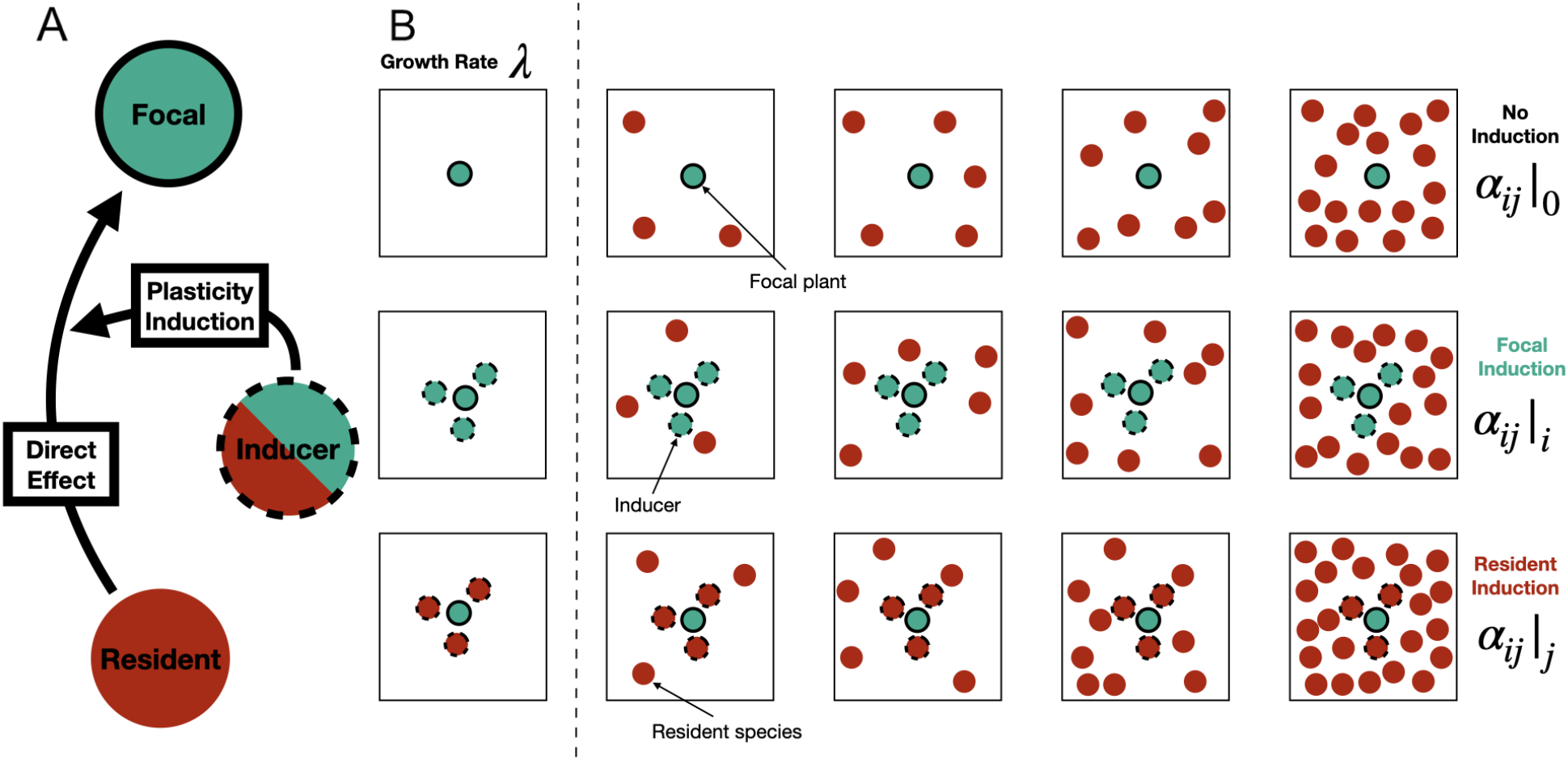
(A) Focal species (in teal with a black line) are temporarily induced by members of either the focal or resident species (the teal and red circle with a dashed outline). We then measure how the induction of plasticity in the focal species modifies the direct effect that the resident species (in red without an outline) has on the focal. (B) To the left of the dashed line, each focal species (teal with solid outlines) is grown with either no inducers (the top row), focal inducers (the middle row with teal and dashed outlined circles) and resident inducers (the bottom row with red and dashed outlined circles). These plots allow inference of the low density growth rate *λ*_*i*_ as well as estimates as the direct competitive effects of the temporary inducers. To the right of the dashed line, other plots vary in the density of the resident species (red circles without outlines), either with no, focal or resident inducers. Note that our design permits the measurement of intraspecific interactions with and without induction when the resident and focal species are the same.

## Materials and Methods

We conducted a field experiment with annual plants designed to quantify how plasticity influences species interactions and ultimately their potential for coexistence. To do so, we designed the study to inform a mathematical model of competition that can predict coexistence outcomes based on the per-capita effects of each species on one another.

### Study site and experimental design

Our study was conducted on four competing annual plant species characteristic of dry calcareous soils in field margins in central Europe – *Buglossoides arvensis, Centaurea cyanus, Papaver rhoeas* and *Sinapis arvensis*. With historically less intensive agriculture, these species took advantage of field habitats in the period after the grain was harvested, but are now relegated to field margins and low quality soils (see Johnson et al. [57] for more information on the system and site). The experiment initially included a fifth species, and its pairwise interactions, but it had very poor germination rates and was thus excluded from the analysis. Seeds for the experiment were collected from monoculture experimental gardens initially established from seeds collected from field edges in Switzerland. In this prior experiment, we obtained estimates of the seed germination rate in these plots by counting the number of germinants in an area of known seed addition, along with the annual seed survival rate following standard protocols (Levine and HilleRisLambers [58]). These rates are reported in Table 2 of the Supplementary Information and were used as parameters in the mathematical framework in Box 1.

Thirty 1.5m by 1.5m plots were established at ETH Zürich’s Research Station for Plant Sciences in Lindau Switzerland in an agricultural landscape. Each plot was surrounded by a snail fence of galvaized metal, separated by 0.75m from neighboring plots, and had soil replaced to 0.5m depth with calcareous soil matching the normal habitat for the study species. All plots were subdivided into two 1.5 by 0.75m subplots. In winter 2015, each of 50 subplots was randomly assigned to receive seeds of one of the five resident species sown at densities of 1, 2, 3, 4, 5, 6, 7, 8, 9, or 10 g m^*−*2^ except for *P. rhoeas*, which required only half those quantities to generate a density gradient. The remaining 10 subplots receive no resident species. In March 2015, we sowed these resident species into the subplots. Each subplot also included 15 equally spaced focal plants (3 individuals of each of the five species). Focal plant positions were 23 cm apart in one direction and 30cm apart in the other, far enough to minimize interactions between focal plants. We did not sow the resident species with 4cm of each focal plant, leaving room for the inducers, as described in the next section. On June 24th, we measured the densities of resident competitors within a 15cm radius of each focal plant to account for the realized neighbor density. Plots were weeded for non-experimental plants throughout the season.

### Induction and competition

At the time of seed sowing, we also established competition induction treatments within 4 cm of each focal individual. To do so, each focal individual was centered within a plastic cylinder (8cm of diameter and 10cm deep) that was partially buried into the soil to minimize seed movement, and then inducing competitors of either the focal species, the resident species, or none as a control were added (Fig. 1). We added enough seed to generate up to 30 germinants. Inducers were thinned on May 7th to between 5-15 individuals depending on the inducing species (see the Supplementary Information for the actual numbers). Early in the growing season, when plants are small, the competitive neighborhood around each focal plant is also relatively small, so these individuals essentially experience competition only from these inducing individuals. On June 1st, after the inducers had time to impact the traits of the focal plants, we removed the inducing individuals to minimize their direct competitive impact. The focal individuals competed only with the background resident species for the remainder of the growing season.

### Data collection

As the plants began to senesce in August, fruits of every focal individual in the different induction and competition treatments were counted. To convert fruit numbers to viable seed numbers, we collected around 50 mature fruits from background resident individuals of each species across the different treatments and then determined the number of seeds per fruit. We then tested a sample of these seeds for viability using standard techniques as in Levine and HilleRisLambers [58]. We also measured a range of plant traits. In early June, soon after the inducers had been removed, we recorded the long axis, short axis, and longest leaf length for five plants from each focal species by induction treatment. We also measured the height of all focal plants in the experiment. A canopy shape measure was calculated as the height divided by the average of the long and short axis.

### Statistical approach

Across species pairs and induction treatments, we fit the Beverton-Holt model to the decline in fecundity of the focal plants of a given species with increasing resident density. Specifically, we fit the following functional form for how fecundity depends on competitor density:

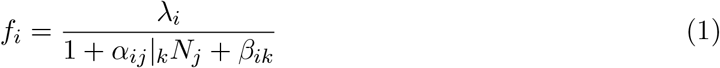

where *f*_*i*_ is the fecundity of an individual of species *i* and *λ*_*i*_ is the seed production in the absence of neighbors for species *i. α*_*ij*_|_*k*_ denotes the per capita competitive effect of species *j* on species *i* when species *i* is induced by species *k*. We let *k* = 0 when the focal species has not experienced induction. Last, *β*_*ik*_ represents both the unavoidable direct competitive effect of the inducing species *k* on focal species *i*. When there is no induction, there is no *β*_*ik*_ term in Equation 1.

#### BOX 1

**Model structure, coexistence theory and phenotypic plasticity**

In the Beverton-Holt model with a seed bank, the dynamics of species *i* are given by

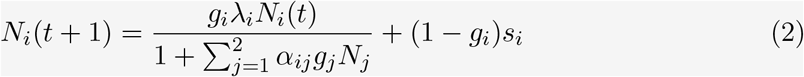

where *N*_*i*_(*t*) is the abundance of species *i* at year *t, λ*_*i*_ is the seed production of species in the absence of neighbors *i, g*_*i*_ is the fraction of seeds that germinate in a given year, *s*_*i*_ is the fraction of seeds that survive a year in the seed bank and *α*_*ij*_ denotes the per-capita competitive effect of species *j* on species *i*.

When species plastically respond to neighbors in ways that affect their interactions, we assume that the *α*_*ij*_ parameters depend on the community composition. Specifically, we denote by *α*_*ij*_|_*k*_ the competitive effect of species *j* on species *i* when species *i* has plastically responded to competition from species *k*. In the theoretical framework of Yamamichi et al. [42], the invasion growth rates can be computed using the interaction parameters where the invading species has plastically responded to a common resident and the resident has plastically responded to itself. The invasion growth rate of species *i* becomes

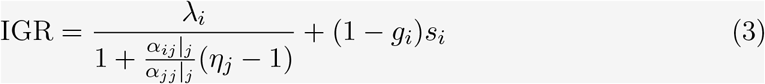

where 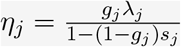. When the invasion growth rates of species *i* and species *j* are both greater than 1, then they can mutually invade one another and are predicted to coexist [37, 42, 59].

The invasion growth rate can be decomposed into a stabilizing niche difference that describes the average ability of species to recover when rare and a fitness difference that describes the average advantage that one species has over the other. While the standard formula for the niche difference between two competitors is 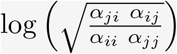, Yamamichi et al. [42], show that when species interaction coefficients are functions of their competitive environment, the stabilizing niche difference can be written as follows:

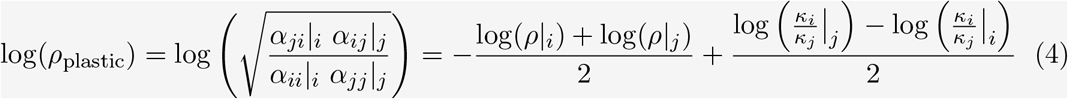

where we have decomposed the niche difference into two useful quantities. In Equation 4, 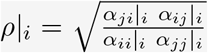 is the niche overlap evaluated when species *i* is common and 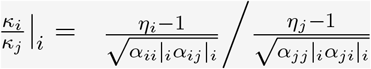 is the fitness difference between species *i* and species *j* when species *j* is common. On the right-hand side of Equation 4, the first term involving only niche differences measures the average niche difference when either species *i* or species *j* is common, while the second term compares the fitness difference between species *i* and species *j* when species *j* is common to when species *i* is common. Intuitively, this first niche difference term describes the ecological niche difference averaged over the plastic responses to the two extreme competitive environments, while the second term measures the benefit to rare species as a result of plasticity. In the context of rapid evolution, these terms correspond to character displacement [43, 44, 45] and the evolution of competitive ability [46, 47, 48, 49, 50] respectively, so we interpret them here as analogous quantities but where trait change is driven by phenotypic plasticity. Last, the fitness difference accounting for the impact of plasticity is

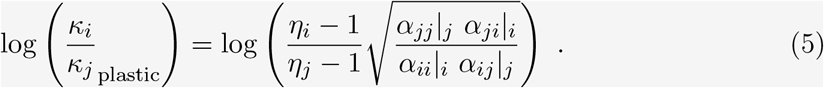

We fit all the models following a Bayesian approach. To construct priors for our fit, we first infer the intrinsic fecundity (*λ*_*i*_) from focal plants in treatments without resident competitors or inducers. We then fit the induction terms (*β*_*ik*_) from focal plants growing with and without inducers, but never with resident competitors. We use the statistics of the inferred distributions of these two parameters (*λ*_*i*_ and *β*_*ik*_) to parameterize normal priors in the fit of the full model. We reduce the variance of these prior distributions to be 10% of the variance of the inferred distributions, so the full model is constrained to produce the growth rates and induction terms that were inferred without competition. We show that our qualitative results are robust to different choices for the variance of the priors in the Supplementary Information. We use weakly informative priors to fit the competition coefficients that are constrained to be positive.

We used the brms package [60] in R version 4.1.2 [61] and assessed mixing and convergence using the R-hat criterion. We ran four chains per model for 10000 total iterations and discarded the first half as warm-up. In the main text, we present results from fitting the Beverton-Holt model because it fit the data well compared to the other models and produced biologically reasonable predictions. Nevertheless, in the Supplementary Information, we consider a suite of different models and compared their fits using leave-one-out cross validation and the widely applicable information criterion. Our qualitative results are robust to different model choices (Supplementary Information). We note considerable uncertainty in the inferred effect of *C. cyanus* on *P. rhoeas* because *C. cyanus* produced very few fruits when induced by either species. Last, we fit linear models for how plant height, plant shape and leaf area responded to the identity of the inducers also following a Bayesian approach. We used uninformative priors, modeled plant height and shape as normally distributed and leaf area as log normally distributed based on leave-one-out cross-validation. We ran four chains per model for 2000 total iterations and discarded the first half as warm-up.

### Coexistence metrics

Although we removed the inducers early in the season, some competitive impact on the focal species is inevitable. To examine how plasticity affects competitive outcomes without this experimental artifact, we fit and then remove the *β*_*ik*_ terms that measure these direct competitive effects. As a result, differences between *α*_*ij*_|_0_ and *α*_*ij*_|_*i*_ or *α*_*ij*_|_*j*_ represent the changes in the strength of species interactions over and above simple reductions in fecundity by the inducers. Note that because our experiments could not manipulate plasticity in the resident background competitor individuals (those individuals can always affect each other’s growth and form), even the “no induction” estimates of interaction coefficients implicitly include the consequences of plastic responses of resident individuals to one another.

We measured the cumulative changes that a given species *i* induces in other species interactions by adding the log ratios of competitive coefficients induced versus not induced by species *i* (ie, 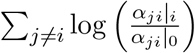 and call this quantity the induction effect. Conversely, we measured the average response of a given species *i* to induction by all other species by summing the log ratios of competitive effects on it when induced versus not induced by the hetero-specific competitor: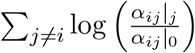. We call this quantity the induction response. Intuitively, these quantities measure how a given focal species effects others through induction and responds to induction from other species.

We used the fitted interaction coefficients and low density fecundity measures, along with the germination and seed survival rates from prior experiments to parameterize the model in Equation 2, Box 1. This allowed us to calculate invasion growth rates with and without induction. In recent work, Yamamichi et al. [42] showed how the invasion growth rates, and niche and fitness differences can provide insight into the dynamics of competing species when the interaction strengths change with neighbors as occurs with plasticity. The key insight is that the interactions relevant for the invasion growth rate and thus coexistence are those where each species’ interactions have adjusted to whichever competitor is common (see Box 1 for a more detailed description). We computed the standard ecological niche differences without induction (Box 1) and the “plastic” niche differences derived in Yamamichi et al. [42] using our inferred interactions and vital rates. Moreover, following methods in Box 1, we decomposed this “plastic” niche difference into a component that captures the ecological niche separation due to plasticity and a separate component that describes the competitive benefit to rare species due to its plastic response (Equation 4 from Box 1). Last we calculated the fitness difference accounting for the impact of plasticity (Equation 5 from Box 1). Because we did not measure how intraspecific interactions changed when induced by a heterospecifc competitor (ie. terms of the form *α*_*ii*_|_*j*_ in Equation 3), we substituted in the intraspecific interactions without induction.

## Results

Plasticity induced by competitors early in life caused plants to be more sensitive to subsequent competition, and interspecific competition in particular (Fig. 2). The exact changes in interaction strength varied depending on the identity of the interacting species. For example, the effect of *S. arvensis* on *B. arvensis* clearly became more competitive when *B. arvensis* responded plastically to competition (regardless of the inducer’s identity), while the reciprocal effect of *B. arvensis* on *S. arvensis* was unchanged (Fig. 2). Plastic responses disproportionately made interspecific interactions more competitive rather than intraspecific interactions, even after accounting for the direct competitive impact of the temporary inducers. Across species, 77.8% of the draws from the posterior distribution indicated more competitive interspecific interactions due to plasticity induced by the focal species compared to 56.7% for intraspecific interactions. **Overall, our first result is that plasticity generally produced more competitive interspecific interactions**.

**Figure 2:**
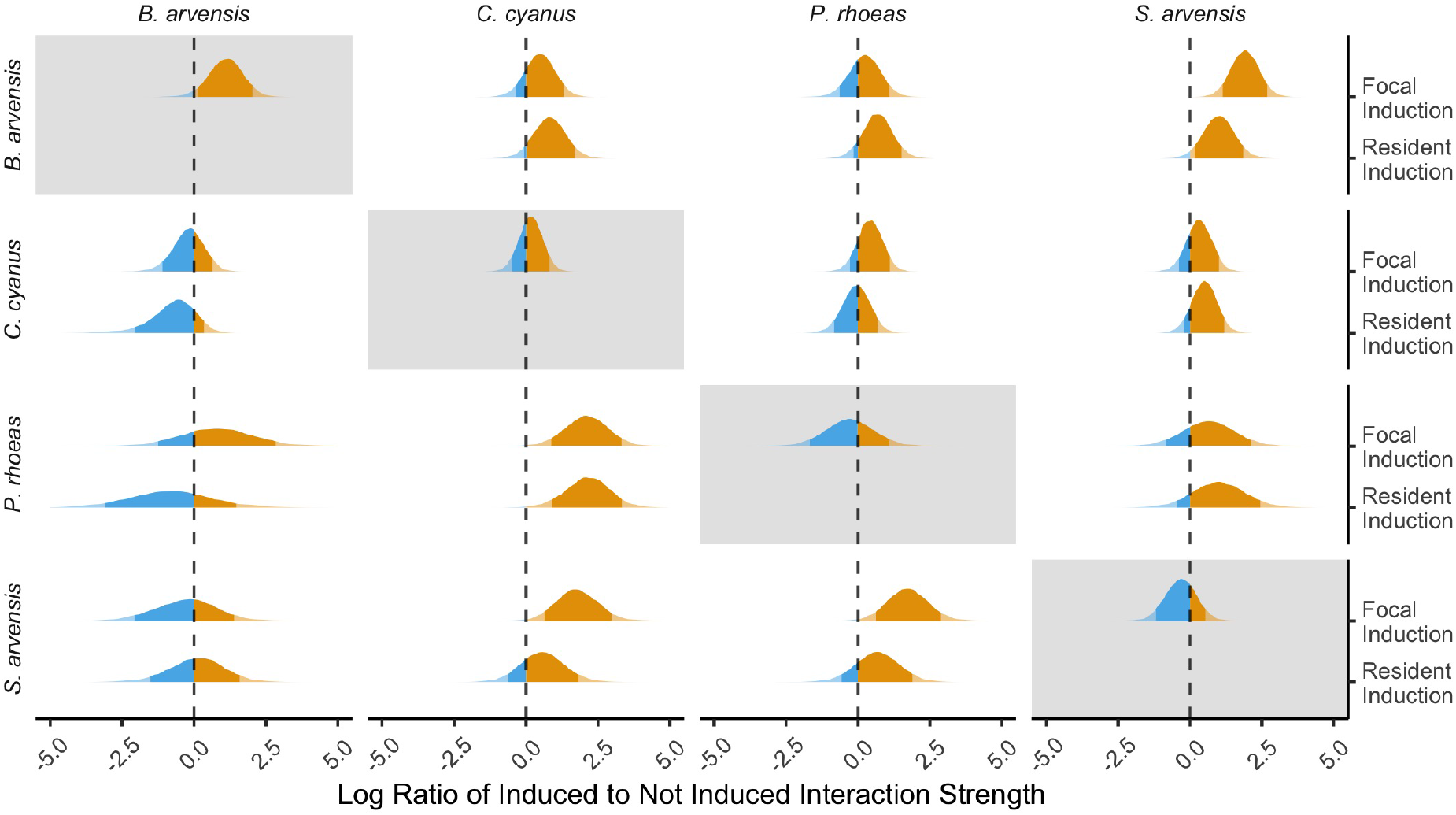
The densities of the posterior distribution of the log ratios of induced to not induced interaction coefficients. The rows display the focal species, while the columns display the resident. For each interaction, we plot how focal and resident induction shift the interaction strength. The densities are colored orange when they are positive, indicating stronger competition as a result of plasticity, while blue colors denote weaker competition. Colors are semi-transparent when outside the 89th percentile of the posterior density. Intraspecific interaction coefficients are highlighted with gray backgrounds.

The species in our study varied in their ability to induce changes in the interactions of other species and the overall sensitivity of their interactions to induction (Fig. 3A). *B. arvensis* had a relatively weak induction effect – often actually reducing the competition experienced by the focal species it induced – but experienced stronger competition as a result of plastic responses to competition early in life (Fig. 3). *C. cyanus* exhibited the opposite dependence, as it strongly amplified competition as an inducer and suffered relatively little from being induced. Across species pairs, we find a weak negative relationship between induction effect and induction response even at the level of individual interactions (Fig. 3B). **In summary, our second main result is that species had characteristic induction effects and responses and that these two quantities tended to be inversely related**.

**Figure 3:**
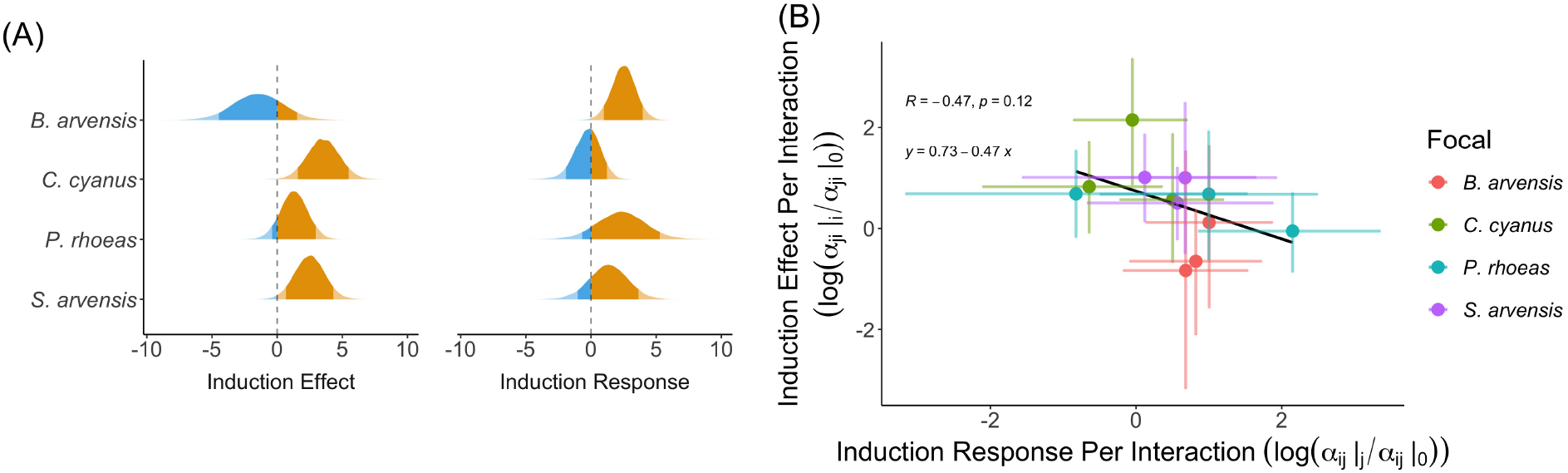
(A) The densities of the posterior distributions for the induction effects and responses (as defined in the Materials and Methods) for each focal species. The densities are colored orange when they are positive, indicating stronger competition as a result of plasticity, while blue colors denote weaker competition. Colors are semi-transparent when outside the 89th percentile of the posterior density. (B) The median induction effect per interaction plotted against the median induction reponse per interaction. Colors denote different focal species and error bars denote the 95% quantiles of the posterior distributions. Line is a linear regression on the medians. The associated correlations, *p*-value and formula are displayed.

Following from our finding that plastic responses to early life competitors enhanced interspecific competition more than intraspecific competition, plasticity tended to reduced the inferred likelihood of one species successfully invading a monoculture of a resident competitor (Fig. 4). In all cases, the predicted distribution of invasion growth rates was lower for at least one of the plastically-responding species. The effect of plasticity on invasion success can be understood using our second main result. With the exception of *B. arvensis*, the intraspecific interaction coefficients did not dramatically shift under induction. On the other hand, the interspecific coefficients tended to become more competitive. As a result, the ratio of interspecific competition to intraspecific regulation increased, disfavoring invasion (Fig. 4). As noted earlier, the only exceptions to this result occurred when *B. arvensis* was the resident – the only case where intraspecific competition did respond considerably to induction.

**Figure 4:**
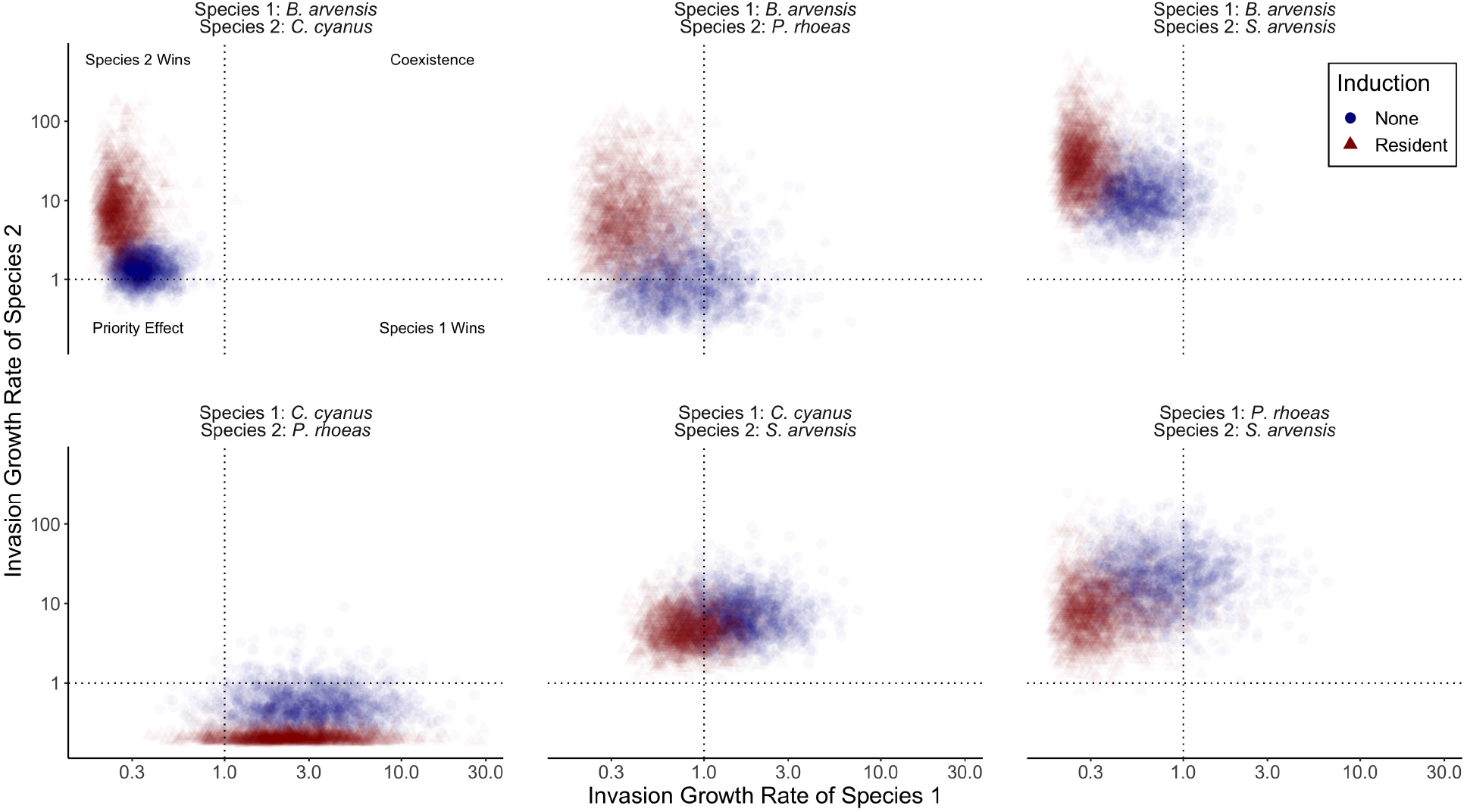
The posterior distributions of the invasion growth rates for each species pair plotted against one another. Points have different colors and shapes for invasion growth rates computed with the resident induced and not induced interaction coefficients. The dashed gray horizontal and vertical lines divide the space into the four possible coexistence outcomes labeled in the top left panel.

Following from the reduced invasion growth rates of plants induced by competitors, plasticity also decreased niche differences between our study species, making coexistence less likely (Fig. 5). More specifically, the niche difference between species *i* and *j*, averaged across the cases where species plastically respond to *i* versus plastically responded to species *j* (first term of the decomposition in Equation 4 in Box 1), tended to be lower than the no induction case for all species pairs not including *B. arvensis* (Fig. 5A). After analyzing the second term in the niche difference decomposition, we found that species dropping to rarity did not appreciably gain competitive ability due to plasticity, and this was consistent across species pairs (Fig. 5B). Last, plasticity had no consistent effect on the fitness differences of the three pairs not involving *B. arvensis*, but tended to decrease fitness differences when *B. arvensis* was present (Fig. 5C). **Overall, our third main result is that plasticity reduced niche differences between species, making coexistence more difficult**.

**Figure 5:**
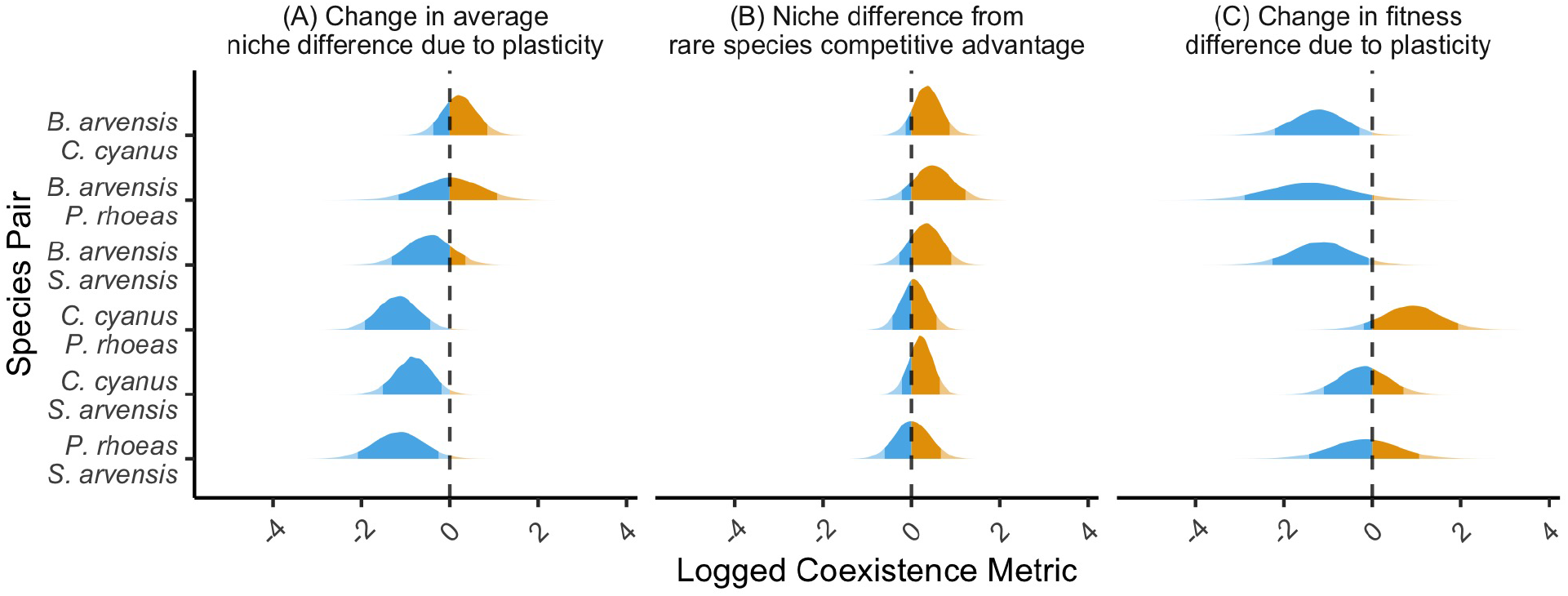
The quantiles and densities of the posterior distribution of the logged values of three coexistence metrics. The y-axis displays the different species pairs. Panels (A) and (B) are the two terms in the decomposition from Equation 4. Panel (A) is the niche difference averaged across cases where either species is common and panel (B) is the niche difference induced by species gaining competitive ability when rare via plasticity. Panel (C) is the change in fitness differences due to plasticity (Equation 5 calculated with and without induction). The dashed vertical lines are located at zero. The densities are colored orange when they are positive, while blue colors indicate negative values. Colors are semi-transparent when outside the 89th percentile of the posterior density.

The traits of the species in our study changed as a result of early-season competition. Growing with nearby competitors early in life tended to make plants shorter or had no effect (Fig. 6A). The only exception was *C. cyanus*, which was taller when induced by members of the same species. Plant shape (the height relative to horizontal extent) did not respond strongly to neighbor induction, except for *B. arvensis* and *C. cyanus* growing more erect when induced by *C. cyanus* (Fig. 6B). The leaves of *B. arvensis* were consistently smaller when experiencing induction by any species (Fig. 6C), which aligns with our previous finding that *B. arvensis* was most sensitive to neighbor induction. **Taken together, our fourth main result is that the traits of some species, and especially *B. arvensis*, one of the more induceable species, noticeably changed as a result of early-season competition**.

**Figure 6:**
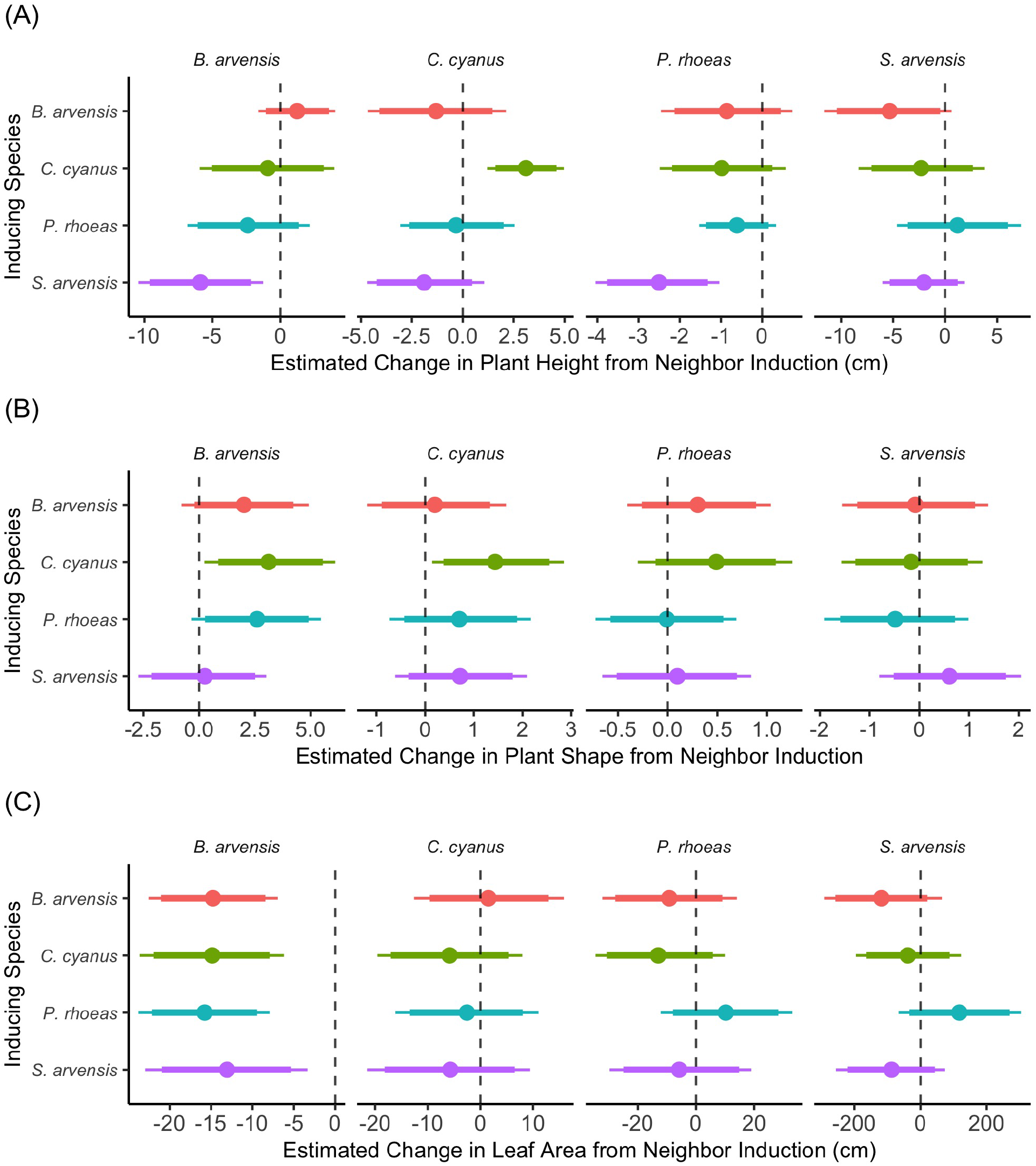
Estimated regression coefficients for how plant height (A), plant shape (B) or leaf area (C) responded to neighbor induction for each inducing species. Points designate the median of the posterior distribution, thick lines show the 89th percentiles and thin lines show the 95th percentiles.

## Discussion

From detailed microbial metabolism [15] to the structure of roots in plants [7], phenotypic plasticity is ubiquitous in natural systems. There is a large body of research demonstrating that competition can be a key mechanism for inducing plastic responses [25, 30, 56, 31], but how the plasticity in turn affects the dynamics of competitors has been poorly resolved. Meanwhile, coexistence research has developed robust methods for inferring species interactions from data but has not considered how plastic responses alter its eventual predictions. As a result, neither of these research areas have fully traced the impact of plasticity from individual traits to species coexistence. Here, we integrated a new experimental design with recent theoretical developments to measure the impact of phenotypic plasticity on species interactions and subsequently coexistence. Specifically, we found that plasticity mostly amplified the competition between our study species, and more so for interspecific than intraspecific interactions (Question 1 from the Introduction). Because of these changes in the interactions, plasticity tended to reduce the predicted probability of coexistence (Question 2 from the Introduction).

We found that the early-season competitive environment affected the eventual interaction strengths, suggesting that competition early in life had a lasting impact on the way the focal species responded to subsequent comnpetition. Because some species traits also shifted in response to early-season competition (Question 3 from the Introduction), we have indirect evidence that changes in height and leaf size may have contributed to changes in the interaction strengths. *B. arvensis*, for example was one of the strongest responders to induction in both its leaf area and interaction coefficients. Still, even at the level of individual traits, those which shifted with early-season competitors were not always associated with individual performance. For example, *C. cyanus* individuals were often taller when exposed to early-season conspecific competitors, possibly reflecting an anticipatory plastic response to avoid competition for light. Yet this species’ intraspecific interaction strength showed no response to induction (Fig. 2). Because we only measured a subset of morphological traits, our analysis of how trait changes affected interaction strengths remains preliminary. Future work should seek to quantify how plasticity in a broader suite of plant traits impacts interaction strengths [62, 63].

Previous studies have put forward the hypothesis that phenotypic plasticity promotes niche differentiation and hence coexistence [24, 33, 34, 35, 64]. These studies have generally drawn these inferences from competition induced differences in dietary preference, phenological timing, or other traits, rather than explicit measurements of how the growth rates and competition strengths change via plasticity. More recent work, however, has manipulated phenotypic plasticity and then determined its effect on the demography of competing species [13, 26, 32, 65].

In these studies as well, plasticity tends to increase the likelihood of coexistence, but not necessarily through niche differentiation [32]. Experimentally parameterized competition models, the approach we took here, allow for a precise definition of niche differences – one of their key strengths – and can therefore evaluate why coexistence outcomes change with plasticity. For example, we found that species pairs not involving *B. arvensis* were less likely to coexist because plasticity generally reduced the niche differences. By contrast, for species pairs involving *B. arvensis*, plasticity tended to reduce fitness differences, rather than increase niche differentiation. Of course, we do not mean to argue that phenotypic plasticity should have predictable effects on coexistence – its effect will depend on the details of the system. Our results do suggest, however, that rigorous and quantitative interrogations of how plasticity affects both inter- and intraspecific interactions are required to unravel the feedback between phenotypic change and community composition.

Our results also illustrate the possibility that the history of interactions a species experiences may influence the strength of the competition affecting its growth in the future. One research direction emerging from our work would involve building mathematical models where each individual has a “memory” (encoded by their morphology) of the competition they have experienced [63, 66, 67, 68, 69, 70]. A similar approach has been proposed in the context of trait-based priority effects, in which the strength of species interactions shifts as a function of trait values which themselves depend on community composition [71, 72].

Although here we have only measured how phenotypic plasticity modifies the interactions between pairs of annual plant species, our results can also be interpreted using theory for higher-order interactions typically applied to systems with three or more competitors. In this framing, we have found that, due to plasticity, each species tends to increase the strength of competition with another species, and more so than it increases competition with individuals of its own species. Interestingly, theoretical work has found that these types of competitive higher-order interactions involving only two species increase the likelihood of coexistence in diverse communities, as long as they are stronger than the higher-order interactions involving three or more species [73, 74]. On the other hand, strongly competitive higher-order interactions among three or more species could erode this benefit to coexistence, so the net effect is difficult to infer.

Because of logistical constraints, we considered only four species and conducted our experiments over just one year. As a result, we cannot evaluate how processes that occur with more species or over longer time scales modify the interactions we have measured. Another limitation of our experiment is that neighbor induction was confined to a short time period early in life. As a result, cross-generational plasticity via maternal effects or epigenetic inheritance could not occur, even though these forms of plastic responses may strongly modify growth rates and interactions [32]. More work is required to determine how the duration and intensity of competition, as well as how different forms of plasticity, influence coexistence. An additional limitation of our study is that we did not experimentally manipulate plasticity in the background competitors harming the focal individuals. Doing so would have been experimentally intractable, but it means that the impact of plasticity in our results is not the complete effect. Despite this concern, it is not clear that plastic responses in the resident competitors differentially affect intra- versus interspecific interactions and therefore may have limited additional effects on coexistence.

Our experimental results reveal the considerable complexity involved in species interactions even at small spatial and temporal scales. An intriguing possibility emerging from our work is that the feedback between phenotypic plasticity and species interactions at these small scales influences broader macroecological patterns. Continued integration of theory and experiment is required first to understand how the feedback between competition and plasticity operates at the level of individual plants and then to propagate these small-scale dynamics upwards to the community level.

## Supplementary Information

The Supplementary Information contains additional figures and descriptions of the methodological approach.

## Acknowledgements

We thank members of the Levin and Levine labs at Princeton University for helpful comments and discussion. We also would like to thank Annabelle Valtat, Ania Donatt, Tabea Kropf, Simon Hart, Myriam von Rütte, Simon Schmid, Andrea Reid, Regina Zaech, and Renato Guidon to for their assistance conducting the experiment or collecting data. This material is based upon work supported by the National Science Foundation Graduate Research Fellowship under Grant No. DGE-2039656 and the High Meadows Environmental Institute at Princeton University through the Mary and Randall Hack ‘69 Graduate Award for Water and the Environment. T.L.G. and J.M.L. acknowledge support from NSF Grant DEB-2022213. M.M.T. acknowledges support from the ETH Zürich Center for Adaptation to Changing Environments and from NSF Grant DEB-1935410.

