## Supplementary Information for "Competitor-induced plasticity modifies the interactions and predicted competitive outcomes between annual plants"

#### Contents

1 Overview 2

2 Qualitative results are robust to variance of priors 2

3 Qualitative results are robust to model choice 3

4 Figures 3

| Name | Functional form |
|---|---|
| Beverton-Holt | $f_i = \frac{s_i \lambda_i}{1+\alpha_{ij}\|_k N_j + \beta_{ik}}$ |
| Law-Watkinson | $f_i = \frac{s_i \lambda_i}{1+N_j^{\alpha_{ij}\|_k} + \beta_{ik}}$ |
| Lotka-Volterra | $f_i = s_i \lambda_i - \alpha_{ij}\|_k N_j - \beta_{ik}$ |
| Ricker | $f_i = s_i \lambda_i e^{-\alpha_{ij}\|_k N_j - \beta_{ik}}$ |

Table 1: The names and functional forms of the models fit to the fecundity data.

### 1 Overview

We fit the four models listed in Table 1 to the fecundity data following a Bayesian approach. As described in the main text, we first fit the $\lambda_i$ values alone and then fit the $\lambda_i$ values and $\beta_{ij}$ values together to arrive at prior distributions for the fit of the full model. In this procedure, we constrained the variance of prior distributions for both the $\lambda_i$ values as well as the $\beta_{ij}$ values to be 10% of the inferred variance. We explore the effects of these choices in the Qualitative results are robust to variance of priors section.

The Lotka-Volterra and Ricker models fit the data worse than the Beverton-Holt and Law-Watkinson models (Fig. A) across all the species, so we did not consider them further. On the other hand, the Law-Watkinson and Beverton-Holt model fit the data similarly. In fact, the Law-Watkinson model fit the fecundity data of some species better than the Beverton-Holt model. As a result, we ensure that our results are qualitatively robust to the choice of model in the Qualitative results are robust to model choice section.

The code used to analyze the data is available at https://github.com/theogibbs/InteractionModification. The GitHub also provides fecundity data (including the number of inducers per focal) and trait data.

### 2 Qualitative results are robust to variance of priors

Priors for $\lambda_i$ with the smallest variance produced the best fits to the data (Fig. A) so we used that approach in the main text. Visualizations of the best fits for Beverton-Holt and Law-Watkinson models, as well as the raw data itself, are provided in Fig. B and C. Increasing the variance of the $\lambda_i$ priors modifies the eventual inferred parameters, especially for $\lambda_i$ and $\alpha_{ij}|_k$ (Fig. D, E and F). For priors with larger variance, the inferred $\lambda_i$ values are larger, but so is the inferred strength of competition ($\alpha_{ij}|_k$), compensating for these larger growth rates (Fig. G). As a result, the ratios between the competition strengths (Fig. H) and relationship between the the invasion growth rates (Fig. I) across different induction treatments are consistent regardless of the variance in the $\lambda_i$ priors. As a result, the central conclusions presented in the main text are not sensitive to the exact choice of variance in the priors for the growth rates.

As described in the main text, we also constrain the variance in the priors for the $\beta_{ij}$ values. While varying the variance in the initial fit of the $\beta_{ij}$ priors, we always use the smallest variance in the $\lambda_i$ priors. The smallest value of the $\beta_{ij}$ prior variance fit the data marginally better than the largest values, so we use the smallest variances in the fits presented in the main text (Fig. J). However, changing the variance in the $\beta_{ij}$ priors had a minor effect on the inferred parameters (Fig. K). Unsurprisingly, our main results are therefore unchanged by the choice of variance in the $\beta_{ij}$ priors (Fig. L).

#### 3 Qualitative results are robust to model choice

Because the Law-Watkinson and Beverton-Holt models fit the data similarly, we consider the inferred Law-Watkinson parameters more closely than the other models we fit. The inferred growth rates and induction parameters are quite similar between the Law-Watkinson and Beverton-Holt models (Fig. M). Although the competition coefficients are not directly comparable between models because of their differing functional forms, they are still clearly correlated (Fig. N), suggesting that the qualitative biological interpretation of these interactions is not model-dependent. The interaction coefficients tended to become more competitive in the Law-Watkinson model (Fig. O), just as in Fig. 3 of the main text. Similarly, we observed the same induction effect and response trends as in Fig. 4 of the main text using the Law-Watkinson model (Fig. P). Last, the qualitative outcomes of pairwise invasions were consistent with our analysis of the Beverton-Holt model, in which induction reduced the likelihood of successful invasion, except when *B. arvensis* was the resident competitor (Fig. Q).

#### 4 Figures

| Species | Seeds per pod | Viability | Germination | Survival |
|---|---|---|---|---|
| *B. arvensis* | 4 | 0.97513 | 0.301 | 0.2306 |
| *C. cyanus* | 18.5 | 0.7384 | 0.46 | 0.4128 |
| *P. rhoeas* | 591 | 0.9567 | 0.034 | 0.1660 |
| *S. arvensis* | 8.7 | 0.982095 | 0.452 | 0.7741 |

Table 2: Seeds per pod and viability, survival and germination probabilities for all species.

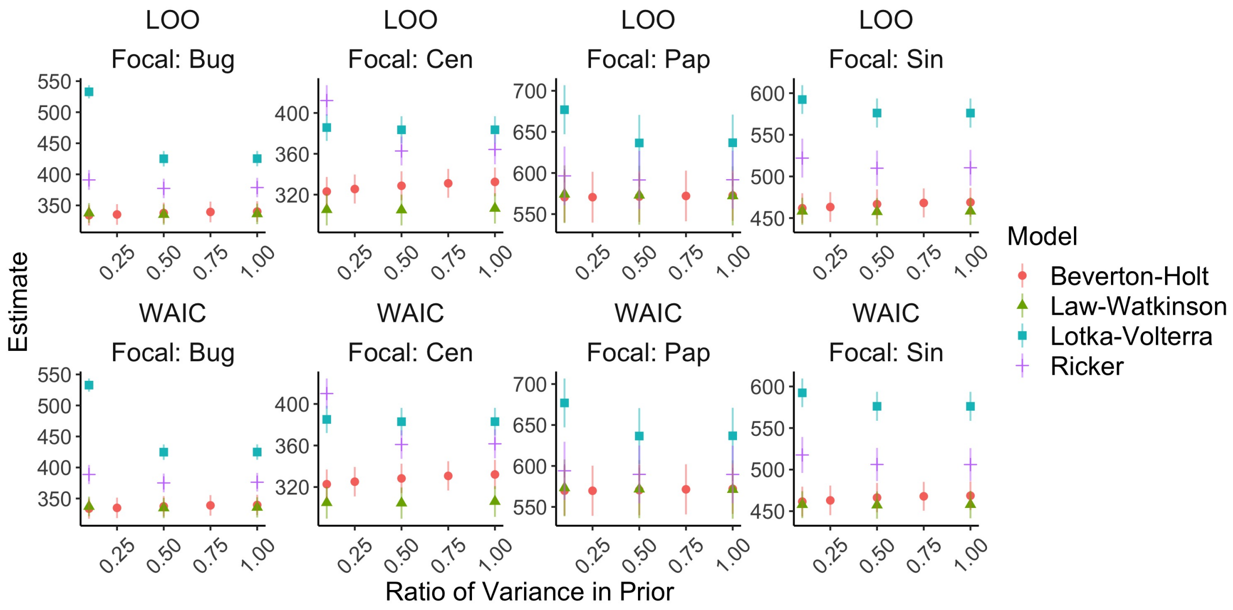


Figure A: Estimates of leave-one-out cross validation (LOO) and the widely applicable information criterion (WAIC) for the different models (colors and shapes) and different choices of the variance in the $\lambda_i$ priors ($x$-axis). Species abbreviations: *B. arvensis* (BUG), *C. cyanus* (CEN), *P. rhoeas* (PAP) and *S. arvensis* (SIN).

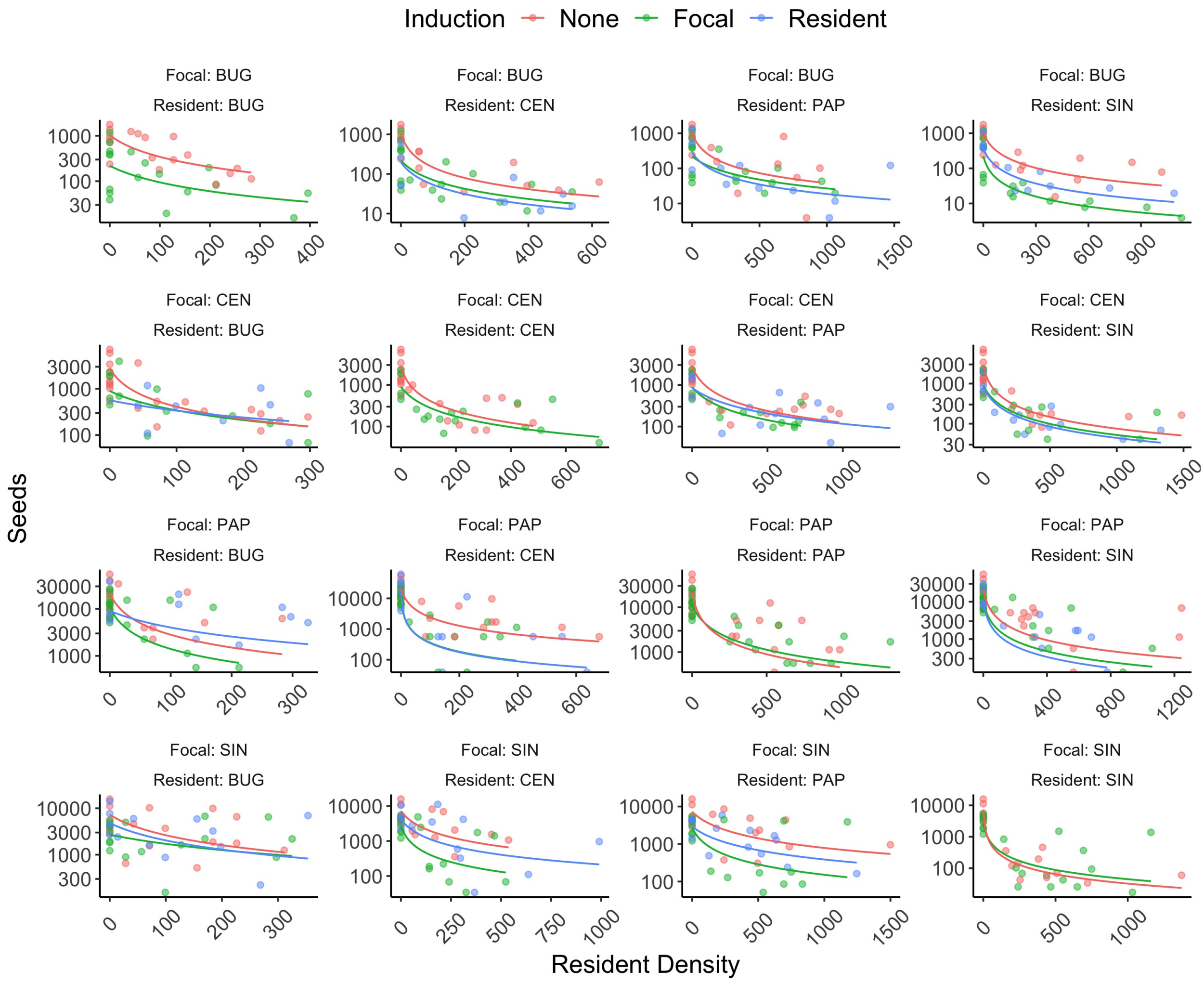


Figure B: Raw fecundity data (points) and the fits of the Beverton-Holt model (lines) across different focal and resident species. Colors designate how the focal species was induced. Species abbreviations: *B. arvensis* (BUG), *C. cyanus* (CEN), *P. rhoeas* (PAP) and *S. arvensis* (SIN).

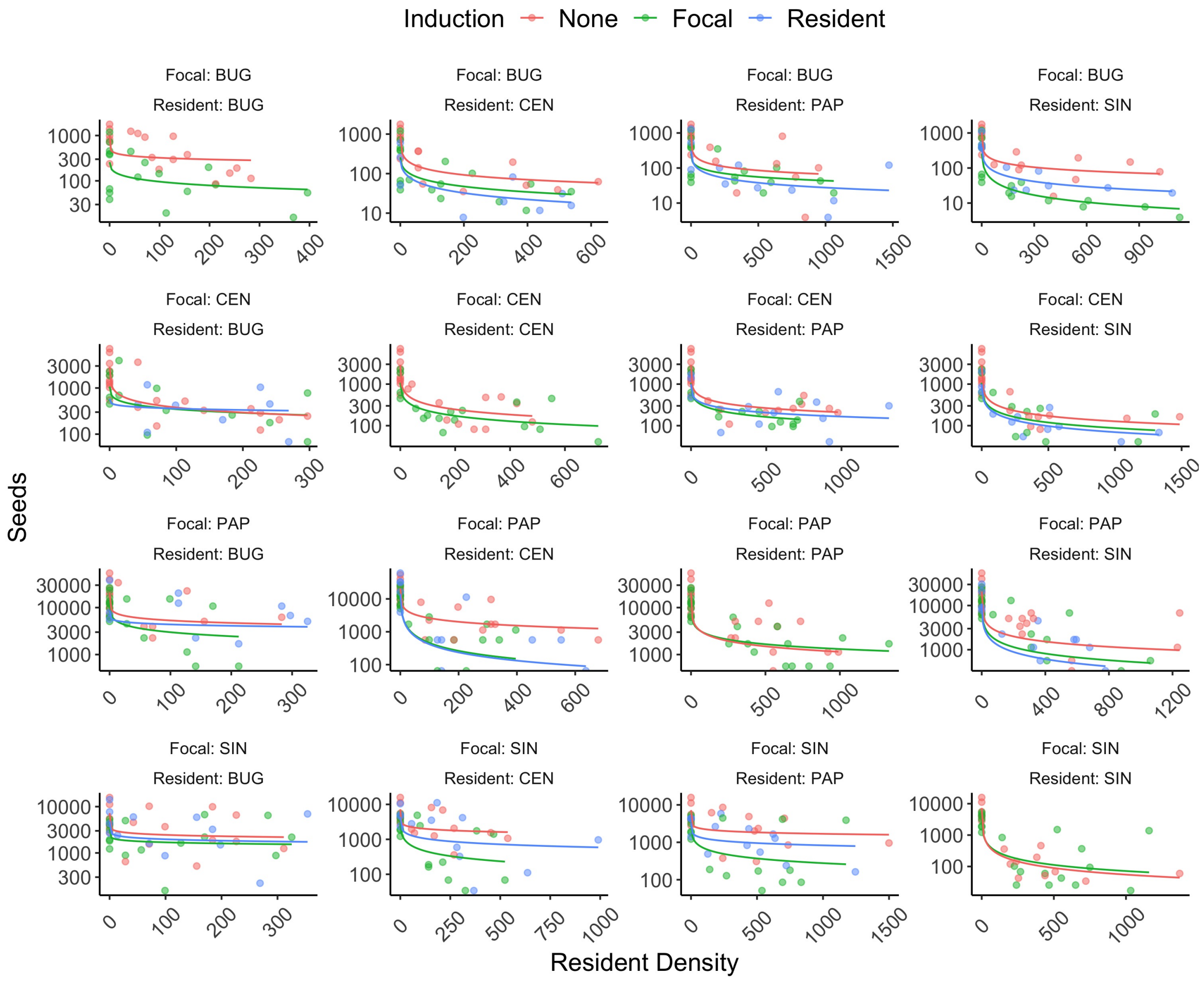


Figure C: Analogous to Fig. B except for the Law-Watkinson model instead of the Beverton-Holt. Species abbreviations: *B. arvensis* (BUG), *C. cyanus* (CEN), *P. rhoeas* (PAP) and *S. arvensis* (SIN).

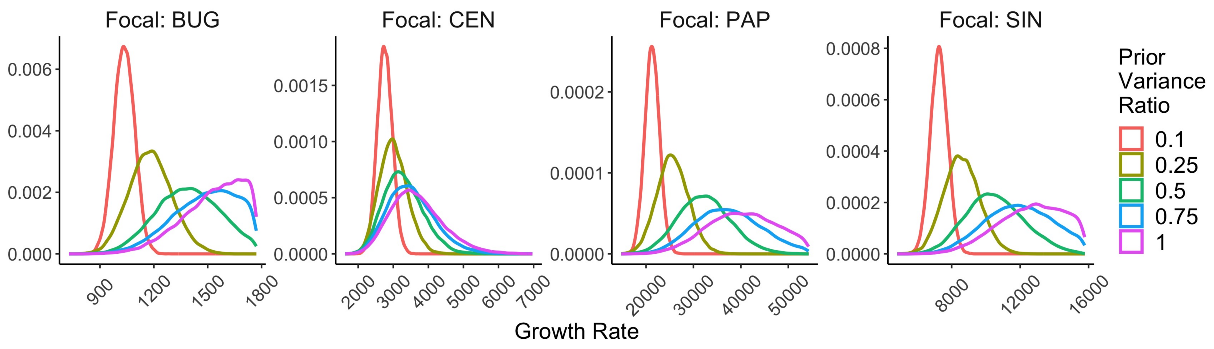


Figure D: The posterior distribution of the $\lambda_i$ parameters in the Beverton-Holt model for the different focal species (panels) and different ratios of the variance in the prior distribution of $\lambda_i$ (colors). Species abbreviations: *B. arvensis* (BUG), *C. cyanus* (CEN), *P. rhoeas* (PAP) and *S. arvensis* (SIN).

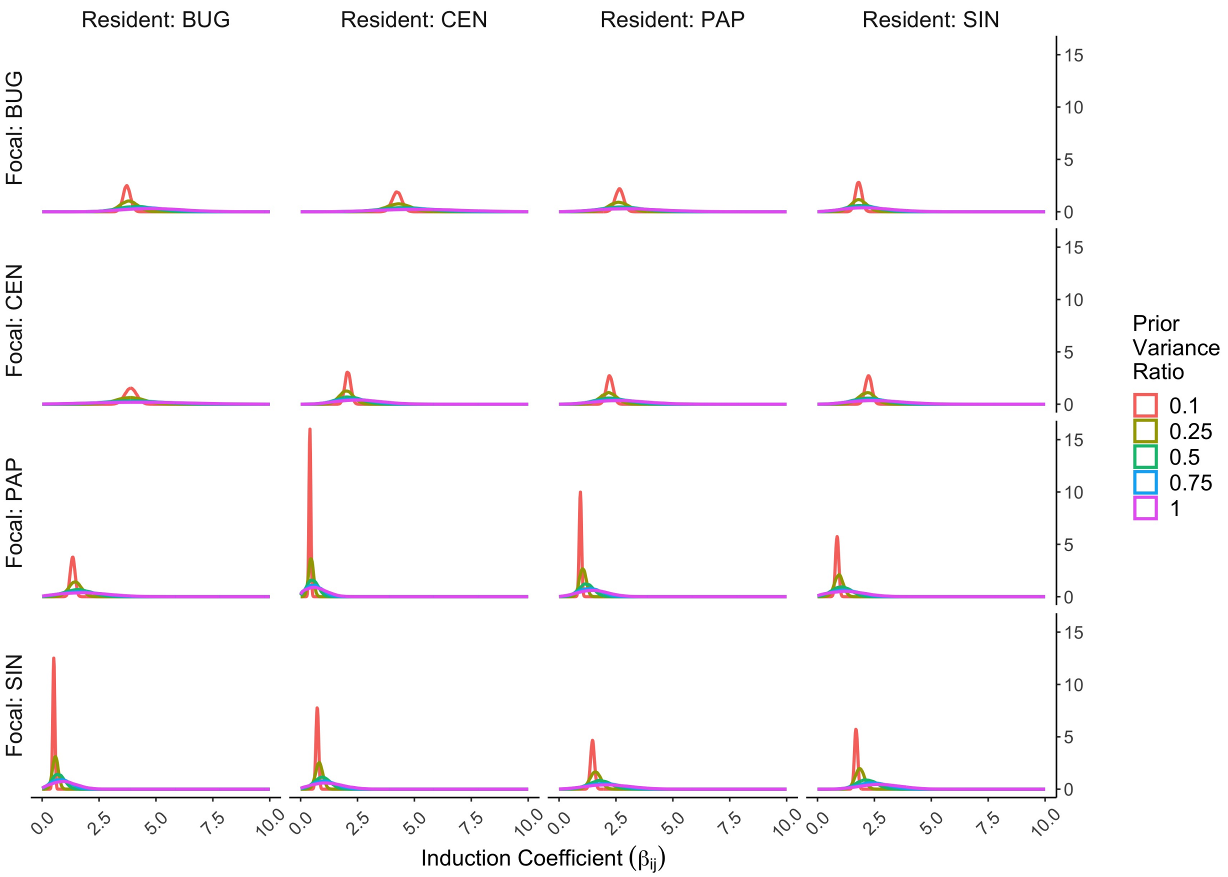


Figure E: The posterior distribution of the $\beta_{ij}$ parameters in the Beverton-Holt model for the different focal species (panels) and different ratios of the variance in the prior distribution of $\lambda_i$ (colors). Species abbreviations: *B. arvensis* (BUG), *C. cyanus* (CEN), *P. rhoeas* (PAP) and *S. arvensis* (SIN).

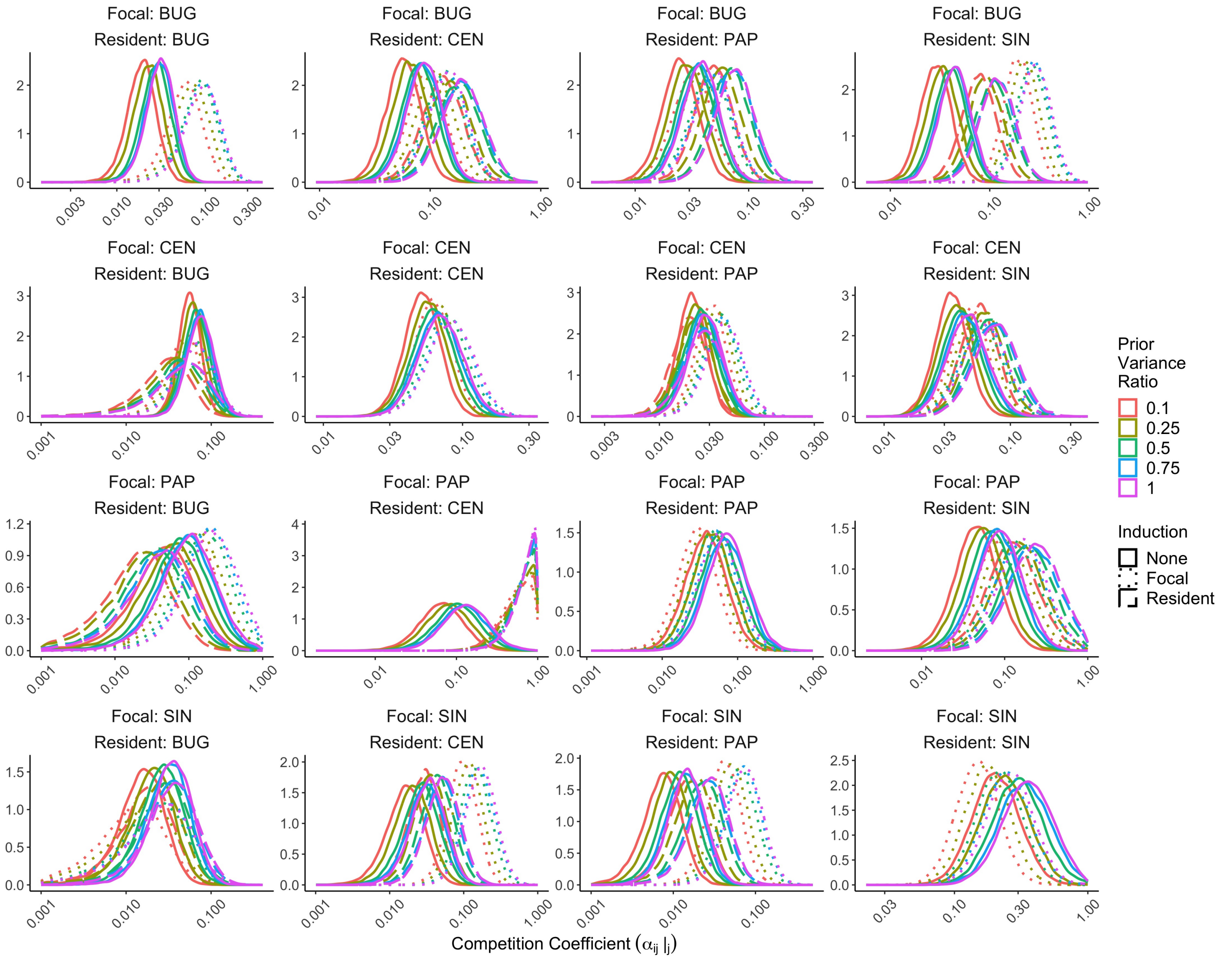


Figure F: The posterior distribution of the $\alpha_{ij}|_k$ parameters in the Beverton-Holt model for the different species pairs (panels), different ratios of the variance in the prior distribution of $\lambda_i$ (colors) and different types of induction (linetypes). Species abbreviations: *B. arvensis* (BUG), *C. cyanus* (CEN), *P. rhoeas* (PAP) and *S. arvensis* (SIN).

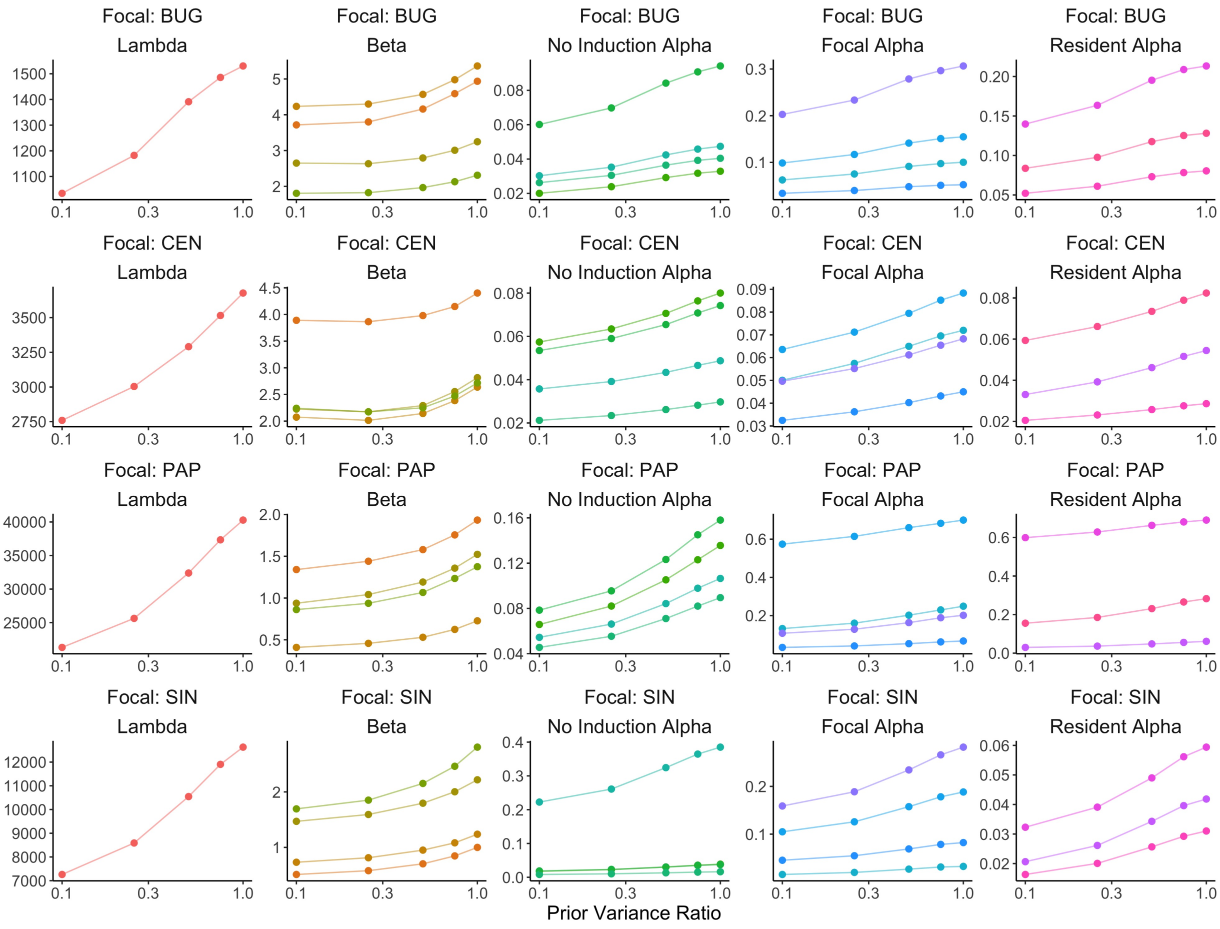


Figure G: The average of the posterior distribution for the different focal species (rows) and model parameters (columns) as a function of the variance in the prior distribution of the $\lambda_i$ values. Colors designate different resident competitors. Species abbreviations: *B. arvensis* (BUG), *C. cyanus* (CEN), *P. rhoeas* (PAP) and *S. arvensis* (SIN).

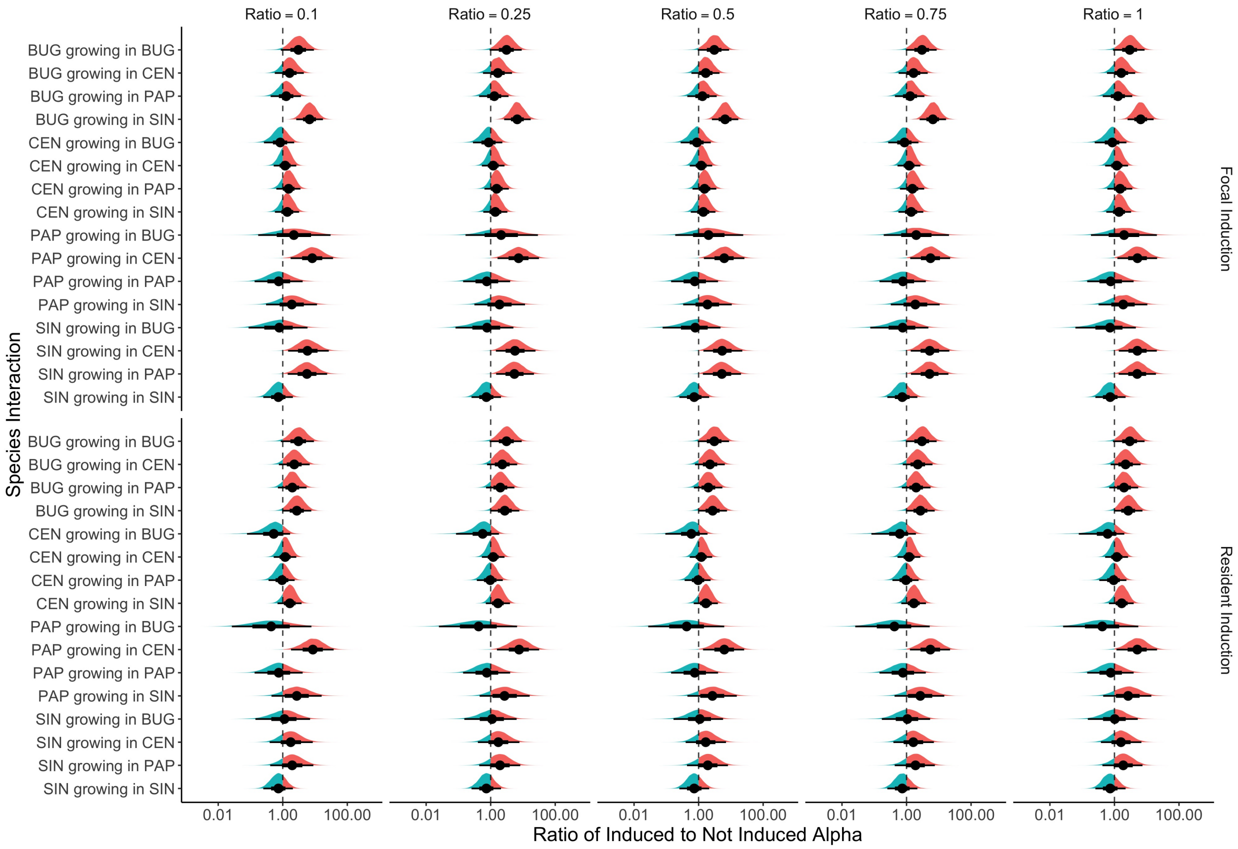


Figure H: Quantiles and densities of the log ratio of induced to not induced competition coefficients. Rows contain different species interactions, rows show focal and resident induction and columns display different variances in the prior distribution for $\lambda_i$. Species abbreviations: *B. arvensis* (BUG), *C. cyanus* (CEN), *P. rhoeas* (PAP) and *S. arvensis* (SIN).

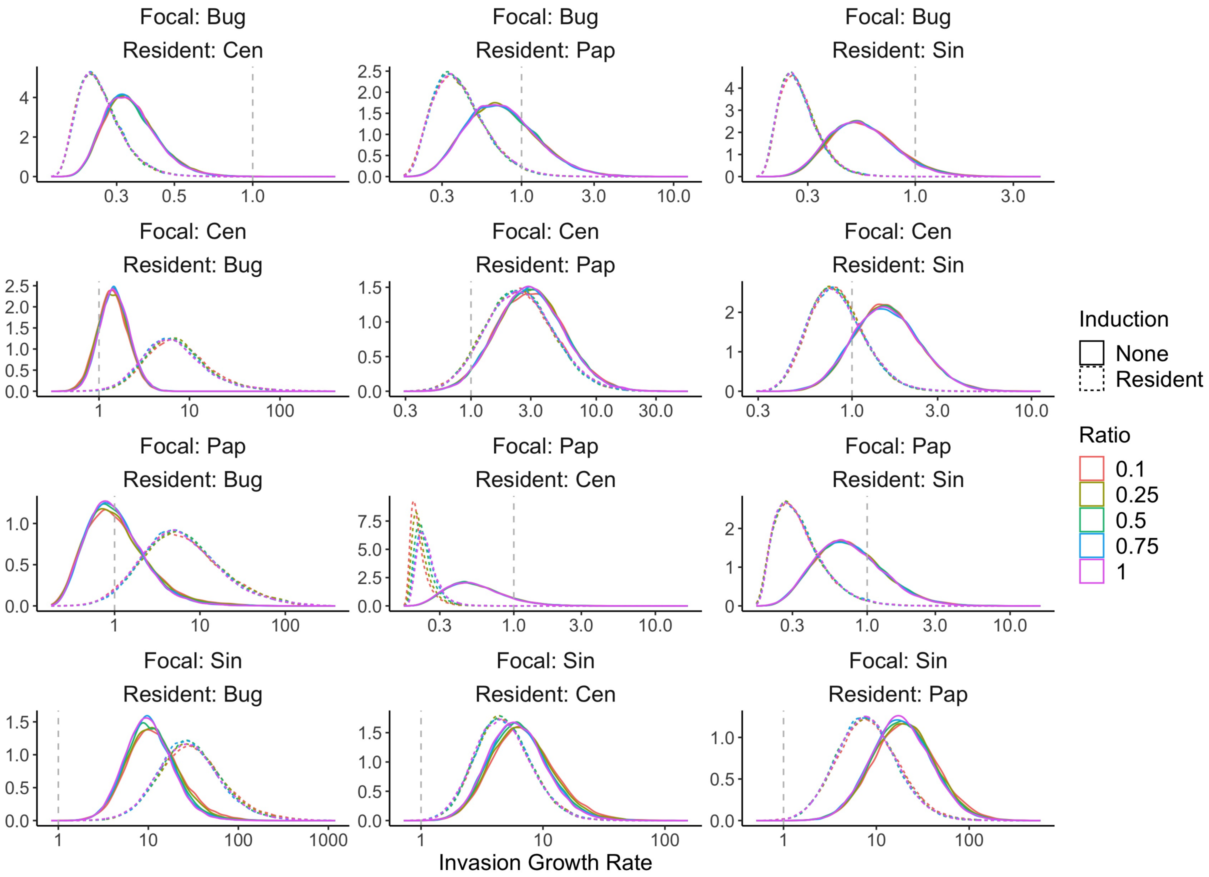


Figure I: Posterior distributions for the invasion growth rates across species pairs (panels), induction treatments (linetype) and the variance in the $\lambda_i$ priors (colors). Species abbreviations: *B. arvensis* (BUG), *C. cyanus* (CEN), *P. rhoeas* (PAP) and *S. arvensis* (SIN).

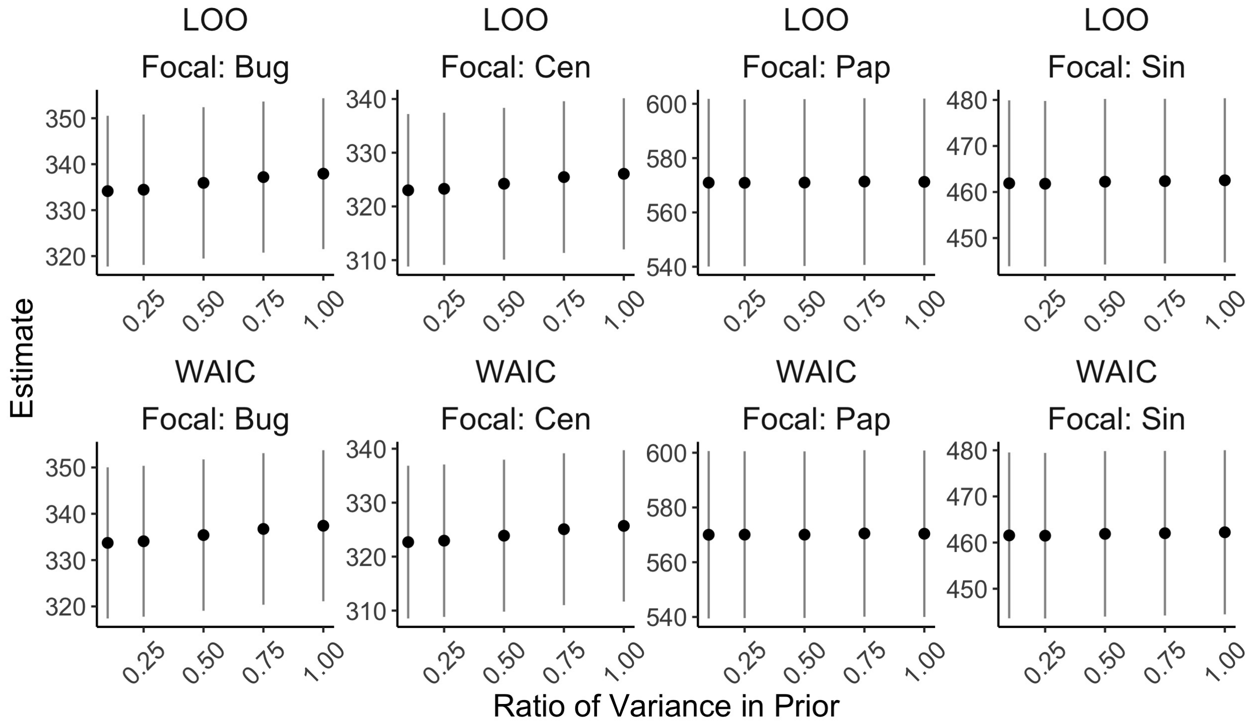


Figure J: Analogous to Fig. A except for the variance in $\beta_{ij}$ priors rather than $\lambda_i$ priors and for only the Beverton-Holt model. Species abbreviations: *B. arvensis* (BUG), *C. cyanus* (CEN), *P. rhoeas* (PAP) and *S. arvensis* (SIN).

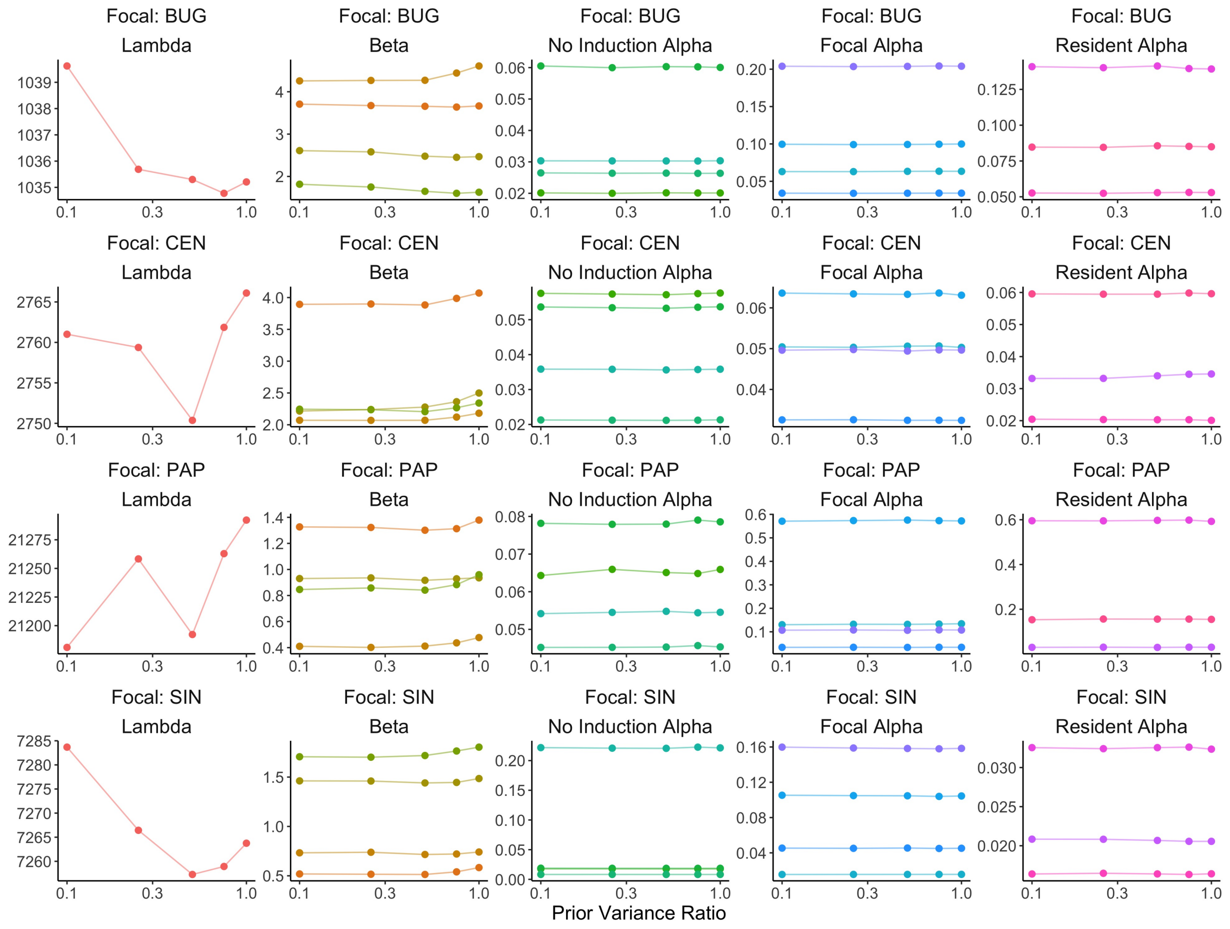


Figure K: Analogous to Fig. G except now for the variance in the $\beta_{ij}$ priors rather than the $\lambda_i$ priors. Species abbreviations: *B. arvensis* (BUG), *C. cyanus* (CEN), *P. rhoeas* (PAP) and *S. arvensis* (SIN).

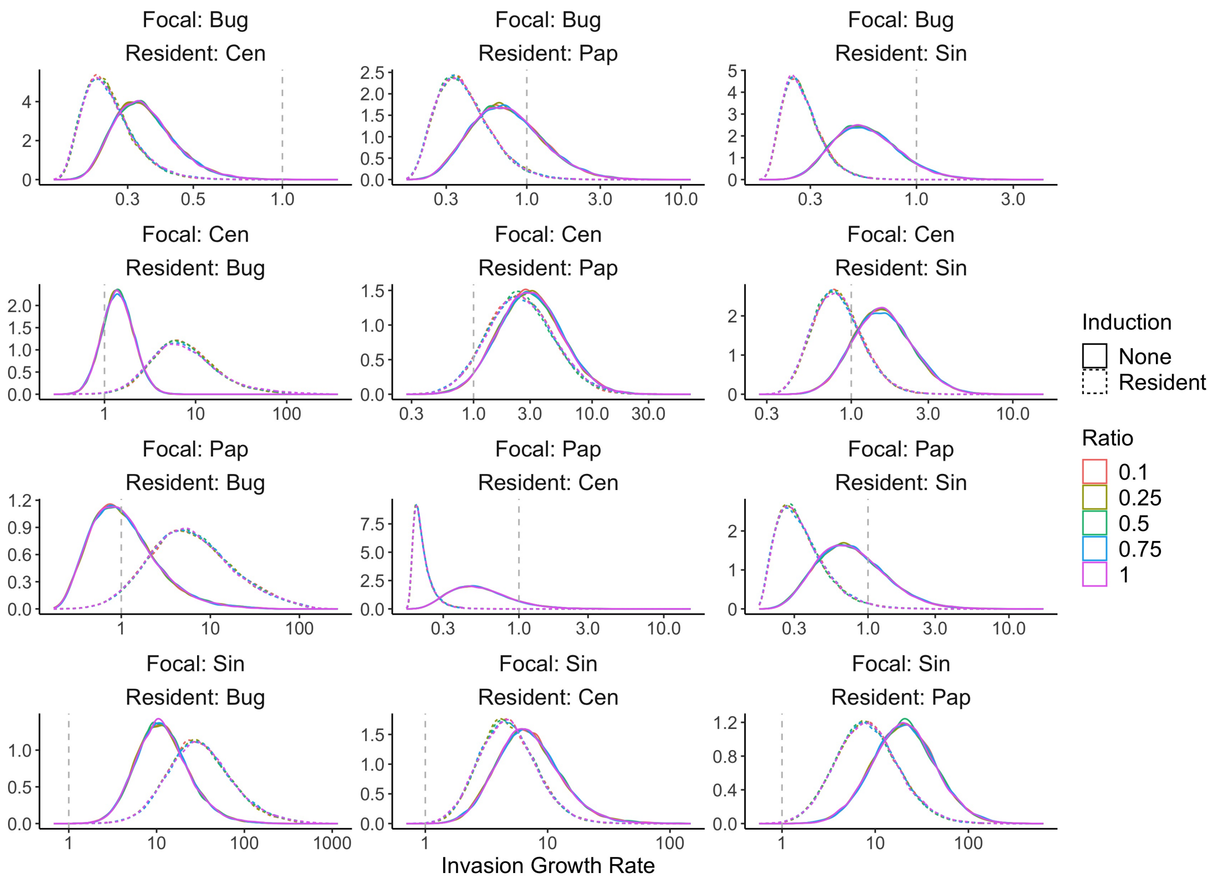


Figure L: Analogous to Fig. I except now for the variance in the $\beta_{ij}$ priors rather than the $\lambda_i$ priors. Species abbreviations: *B. arvensis* (BUG), *C. cyanus* (CEN), *P. rhoeas* (PAP) and *S. arvensis* (SIN).

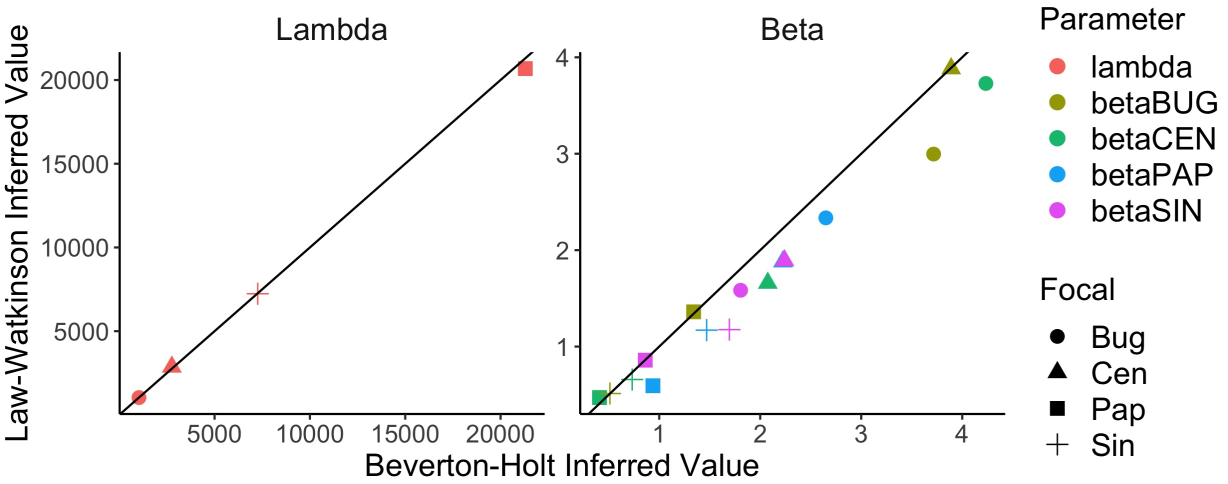


Figure M: The values of the growth rates and induction terms (panels) inferred using the Beverton-Holt ($x$-axis) and Law-Watkinson ($y$-axis) models. Colors show different parameter values and shapes show different focal species. Solid lines show $y = x$. Species abbreviations: *B. arvensis* (BUG), *C. cyanus* (CEN), *P. rhoeas* (PAP) and *S. arvensis* (SIN).

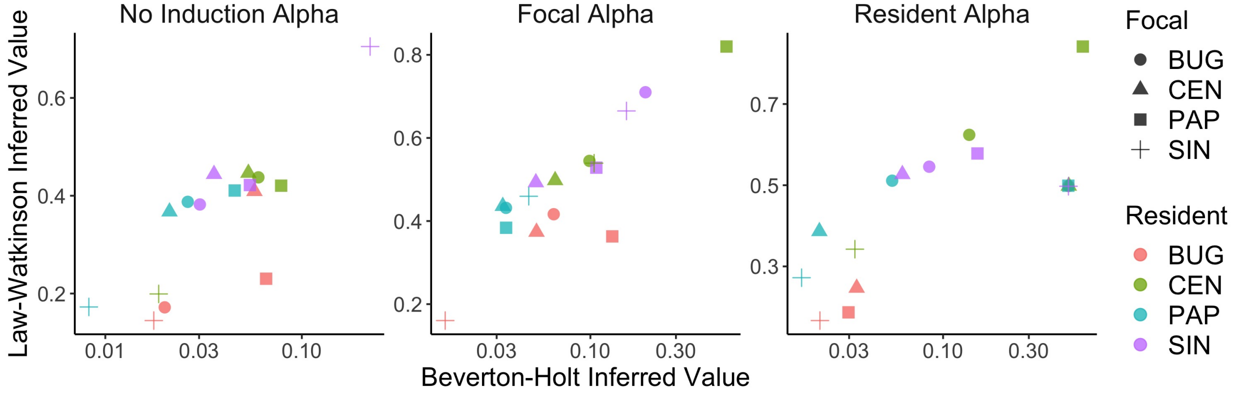


Figure N: The values of the competition coefficients under different induction treatments (panels) inferred using the Beverton-Holt ($x$-axis) and Law-Watkinson ($y$-axis) models. Colors show different resident species and shapes show different focal species. Species abbreviations: *B. arvensis* (BUG), *C. cyanus* (CEN), *P. rhoeas* (PAP) and *S. arvensis* (SIN).

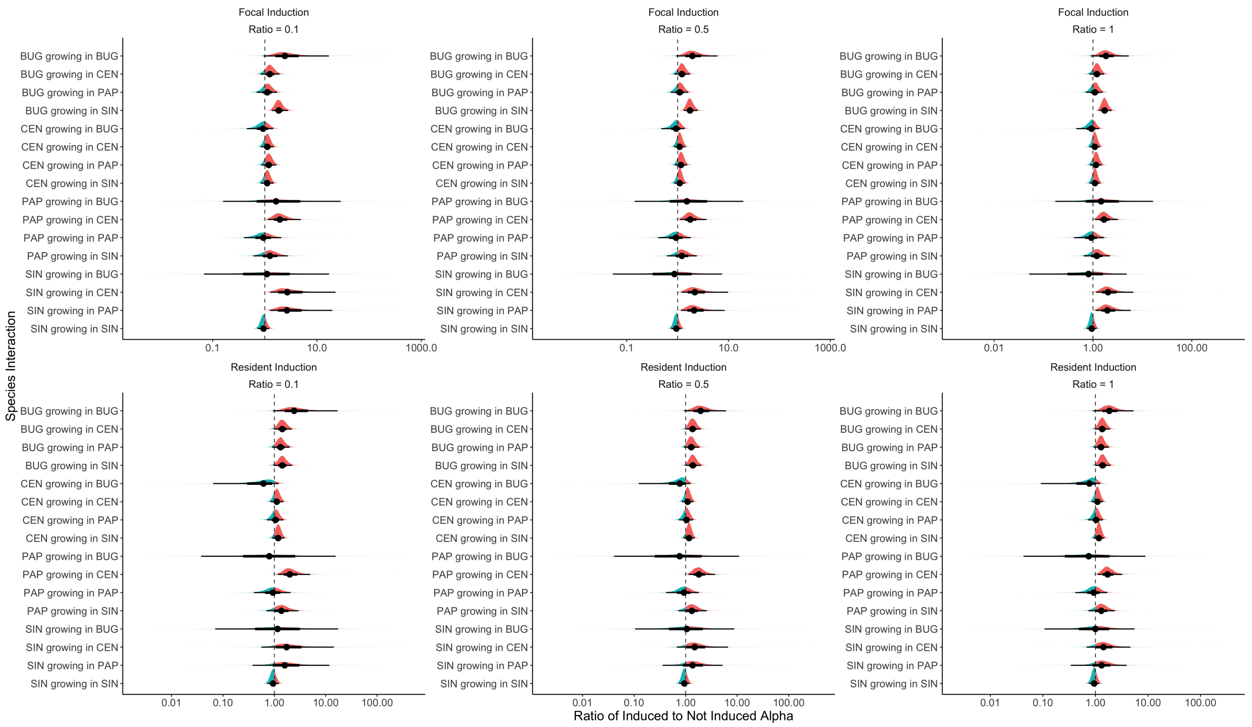


Figure O: Analogous to Fig. H except for the Law-Watkinson model. Different $\lambda_i$ ratios are shown across the columns. Species abbreviations: *B. arvensis* (BUG), *C. cyanus* (CEN), *P. rhoeas* (PAP) and *S. arvensis* (SIN).

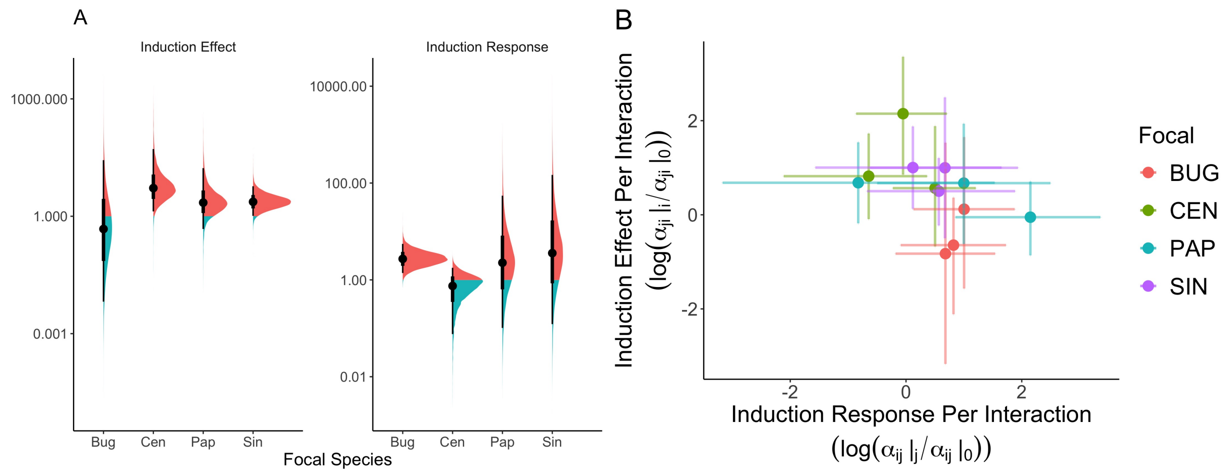


Figure P: Analogous to Fig. 4 in the main text but with parameters inferred from the Law-Watkinson model. Species abbreviations: *B. arvensis* (BUG), *C. cyanus* (CEN), *P. rhoeas* (PAP) and *S. arvensis* (SIN).

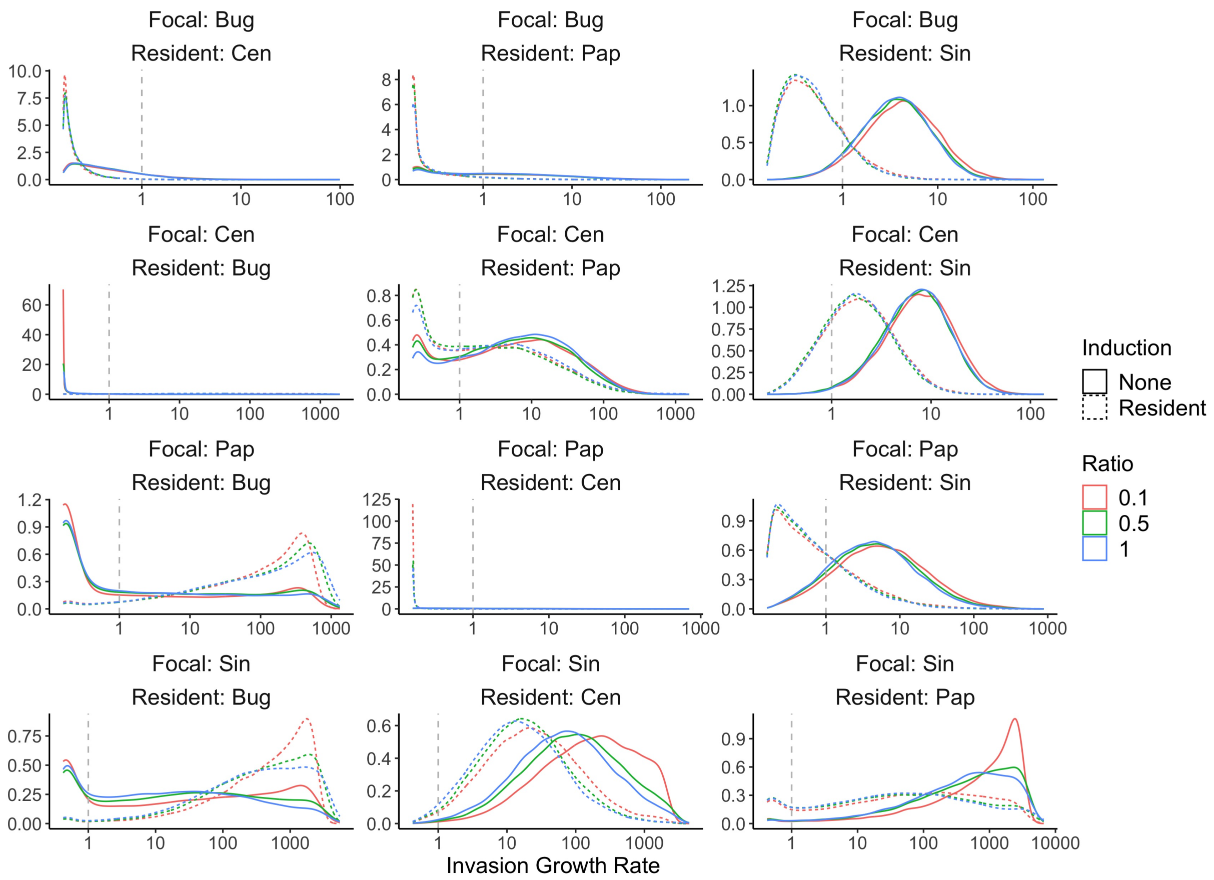


Figure Q: Analogous to Fig. I but using parameters inferred with the Law-Watkinson model. Species abbreviations: *B. arvensis* (BUG), *C. cyanus* (CEN), *P. rhoeas* (PAP) and *S. arvensis* (SIN). The posteriod distributions are multimodal because the inferred competition coefficients produce invasion growth rates over a large range of scales, but there is a minimum invasion growth rate set by the combination of the germination and survival probabilities in the functional form of the model.
